# Accessible chromatin regions and DNA methylation regulate gene expression leading to changes in agronomic traits in *Brassica* allotriploid hybrids

**DOI:** 10.1101/2024.09.18.613586

**Authors:** Chengtao Quan, Qin Zhang, Xiaoni Zhang, Kexin Chai, Guoting Cheng, Chaozhi Ma, Cheng Dai

## Abstract

**Introduction:** Interspecific hybridization is a common method in plant breeding to combine traits from different species, resulting in allopolyploidization and significant genetic and epigenetic changes. However, our understanding of genome-wide chromatin and gene expression dynamics during allopolyploidization remains limited.

**Objectives:** We aimed to explore the relationship and underlying mechanisms between accessible chromatin regions and DNA methylation and gene transcription in genome-wide reorganization after interspecific hybridization.

**Methods:** This study generated two *Brassica* allotriploid hybrids via interspecific hybridization, combining transcriptomics, whole-genome bisulfite sequencing (WGBS) and assay for transposase-accessible chromatin with high throughput sequencing (ATAC-seq), revealing that accessible chromatin regions (ACRs) and DNA methylation regulate gene expression after interspecific hybridization, ultimately influencing the agronomic traits of the hybrids.

**Results:** A total of 234,649 ACRs were identified in the parental lines and hybrids, the hybridization process induces changes in the distribution and abundance of there accessible chromatin regions, particularly in gene regions and their proximity. On average, genes associated with Proximal ACRs were more highly expressed than the genes associated with Distal and Genic ACRs. More than half of novel ACRs drove transgressive gene expression in the hybrids, and the transgressive up-regulated genes showed significant enrichment in metal ion binding, especially magnesium ion, calcium ion, and potassium ion binding. We also identified the *Bna.bZIP11* in the single-parent activation ACR (SPA-ACR), which binds to *BnaA06.UF3GT* to promote anthocyanin accumulation in F_1_ hybrids. Additionally, in F_1_ hybrids, the level of DNA methylation in ACRs was higher compared to gene bodies, and the A-subgenome ACRs were associated with genome dosage rather than DNA methylation.

**Conclusions:** The interplay among DNA methylation, TEs, and sRNA contributes to the dynamic landscape of ACRs during interspecific hybridization, resulting in distinct gene expression patterns on the genome.

**HIGHLIGHTS:**

- The study utilized the accessible chromatin regions (ACR) and DNA methylation to elucidate the mechanism behind gene expression changes following interspecific hybridization.
- Whole-genome recombination after interspecific hybridization leads to the rearrangement of ACR, and novel ACR and single-parent activation ACR regulate the expression of genes.
- DNA methylation plays a role in repressing gene expression within ACRs, and unmethylated ACRs have more transcriptionally active.
- A-subgenome ACRs were associated with genome dosage rather than DNA methylation.

**GRAPHICAL ABSTRACT:** 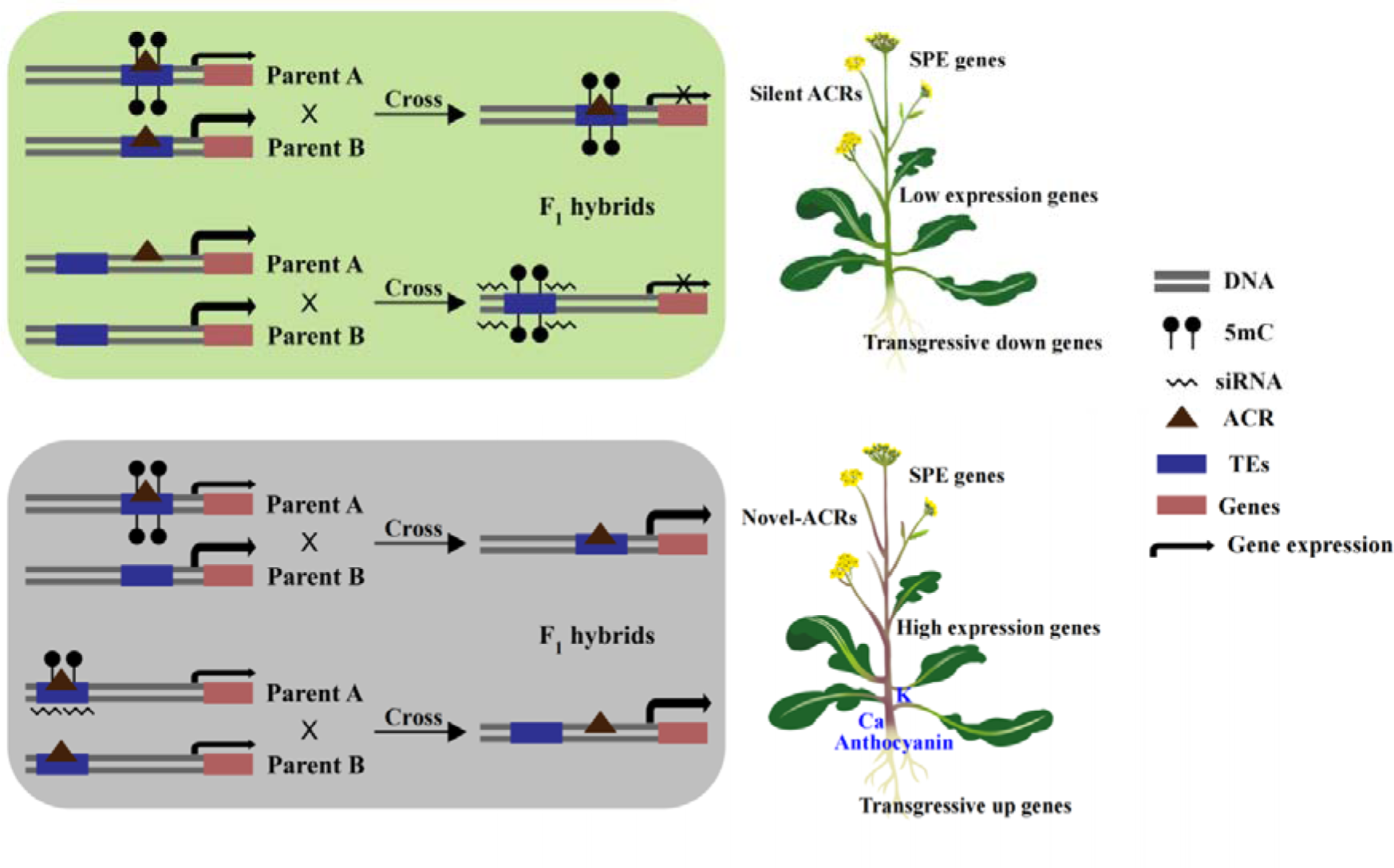

## Introduction

Interspecific hybridization is an essential tool in plant breeding and genetic improvement. It allows the incorporation of desirable traits from different species into a single organism, improving crop quality and productivity [1–4]. This process has revolutionized crop breeding and contributed significantly to the global food supply [5,6]. A major advantage of interspecific hybridization is the generation of hybrids with superior vigor compared to their parents. This vigor arises from interactions between alleles at multiple loci, epistatic interactions, and possibly complementation of deleterious mutations [7]. Emerging evidence suggests that epigenetic factors are crucial for hybrid vigor [2,8]. However, the genetic mechanisms underlying interspecific hybridization are complex and not yet fully understood.

Accessible chromatin regions (ACRs) are typically nucleosome-free or loosely bound to nucleosomes, making them more susceptible to binding by regulatory proteins that affect gene expression. Pinpointing ACRs is critical to understanding *cis*-regulatory elements (CREs) in the genome, which form the intricate transcriptional regulatory networks that control gene expression [9]. In tea plants, hybrids have more ACRs than the parents, which may confer broader and stronger transcriptional activity to genes in the hybrids [10]. The ACRs provide valuable insights into how the physical structure of chromatin influences gene expression and contributes to the superior traits observed in hybrids [9,10]. Thus, uncovering these ACRs and their associations with gene expression profiles may help to unravel the complex regulatory networks behind hybrid vigor.

DNA methylation is a vital process of epigenetic modification in which methyl groups are added to the DNA molecule. It is a critical factor in regulating gene expression without affecting the genetic sequence [11,12]. In *Arabidopsis thaliana*, DNA methylation in all three sequence contexts was observed to impact chromatin accessibility in heterochromatin. Loss of DNA methylation in one or two sequence contexts resulted in the majority of chromatin regions remaining inaccessible. However, when DNA methylation was reduced in all contexts, there was a notable impact on accessibility characteristics, highlighting the key role of DNA methylation in maintaining heterochromatin inaccessibility [13]. Most DNA methylation patterns in plants are generally stable, but DNA methylation in CHG and CHH contexts can be altered following interspecific hybridization [14]. Research indicates that most regions of chromatin accessibility are hypomethylated, and that changes in DNA methylation induced by acclimation may affect chromatin accessibility to influence the expression levels of nearby and distant genes [15,16].

The Brassicaceae family of species is essential in providing vegetable oil, animal feed, and vegetables for human consumption worldwide [17,18]. Interspecific hybridization has been widely used in the *Brassica* genus to combine desirable traits from different species, resulting in new crop varieties with improved agronomic performance [19]. In this study, we have identified epigenetic modifications and changes in chromatin accessibility that contribute to heterotic traits in the interspecific hybridization of *Brassica napus* and *Brassica rapa*. This information can be used to develop breeding programs aimed at improving crop performance.

## Materials and Methods

### Plant material

Two *Brassica napus* cultivars (s*70*, A_s_A_s_C_s_C_s_; *yu25*, A_y_A_y_C_y_C_y_, allotetraploid) were selected as the maternal parent, and *Brassica rapa* cultivar (*B. campestris L. ssp. chinensis var. purpuria* Hort.; A_h_A_h_, diploid) were selected as the paternal parent. The F_1_ allotriploid hybrids were generated by crossing between two different *B. napus* and the *B. rapa* species. All plant materials were grown under the same field conditions located at the Huazhong Agricultural University (30°28ʹN, 114°21ʹW). For sampling, the plant materials were collected at 10.00 – 11.00 am in December (average 8°C). Stem epidermal tissue below the top of the fifth true leaf was collected and immediately frozen in liquid nitrogen. These samples were then used for subsequent ATAC-seq, WGBS, sRNA-seq, and RNA-seq library construction, and for determining ions and metabolites.

### ATAC-seq analysis

The low-quality reads and adapters from the raw ATAC-seq data were filtered and removed using Trimmomatic [20]. The clean data were then aligned to the *B. napus* (Zhongshuang 11, ZS11) reference genome by Bowtie2_v2.5.2 [21,22]. The mapped reads in sam format were converted to bam format using SAMtools_v1.9 [23]. Subsequently, MACS2_v2.2 peak calling software was used to identify ATAC-seq peaks [24]. The overlapping peaks over 50 bp in the biological replicates were considered as ACRs. The genomic distribution of ACRs and associated genes was confirmed using the ChIPseeker tool [25]. Differential binding events were identified using the DiffBind_v6.0 package [26]. The motifs of the ACRs were identified using HOMER [27]. The term “silent ACR” refers to ACRs identified in parents (reads ≥ 5) but not in F_1_ hybrids (reads = 0). On the other hand, the term “novel ACR” refers to ACRs identified in F_1_ hybrids (reads ≥ 5) but not in parents (reads = 0).

### WGBS analysis

We then used BatMeth2-align with default parameters to map the filtered WGBS reads to the *B. napus* (ZS11) genome [28]. The sequences covering 5 or more cytosine sites were set as valid methylation sites. Finally, BatMeth2-Meth2BigWig was used to generate BigWig files to identify and visualize differentially methylated regions (DMRs) in IGV [28]. Only cytosine regions with adjusted *p*-values < 0.05 and DNA methylation differences greater than 0.3, 0.2, and 0.1 (for CG, CHG, and CHH, respectively) were considered DMRs.

### RNA-seq analysis

Trimmomatic was used to remove barcode adaptors and low-quality reads [20]. The filtered reads were then aligned to the *B. napus* (ZS11) reference genome using HISAT2_v2.2.0 with default parameters [29]. The uniquely mapped reads were filtered using SAMtools_v1.9 [23]. Read counting and normalization of transcripts per million mapped reads (FPKM) were performed on BAM files using StringTie_v2.1.4 [30]. Genes with FPKM > 1 were defined as expressed genes. Genes with an adjusted *p*-value < 0.05 found by DESeq2 and a |log_2_fold change| ≥ 2 were assigned as differentially expressed [31].

### sRNA-seq analysis

The raw sequencing reads were trimmed using cutadapt to remove adapters (https://github.com/ marcelm/cutadapt/tree/v3.1). Subsequently, sRNAs between 18 and 30 nt in length were selected and mapped to the *B. napus* (ZS11) genome and defined into sRNA clusters using Shortstack_v3.8.4 [32]. sRNA-mapped reads were normalized to the total cleaned reads for further analysis.

### Elemental mass spectrometry analysis

The concentrations of mineral elements in the stem and epidermis were measured using an inductively coupled plasma-mass spectrometer (ICP-MS) (Perkin Elmer, NexION 300D, Shropshire, UK). The replicate samples for each line were pooled and then subjected to oven-drying at 80 °C for a minimum of 72 h, followed by grinding. Approximately 0.2 g of the dried ground powder was placed in a PTFE digestion tube along with 6 mL of concentrated nitric acid, and the tube was tightly sealed and processed in a closed vessel acid digestion microwave (MARSXpress; CEM Corporation, Matthews, NC, USA). After digestion, each digested sample was diluted to 10 mL with deionized water, and elemental analysis was performed using an ICP-MS in the standard mode, monitoring 15 elements.

### Measurement of metabolites

The measurement of soluble sugars, soluble proteins, total phenolics, total flavonoids, total anthocyanins, and proanthocyanidins in the stem epidermal tissue was performed as described in previous studies [33]. All samples were quantified in triplicate in three independent biological replicates.

### Electrophoretic mobility shift assay

The full-length CDS of *BnaA03.bZIP11* and *BnaC03.bZIP11* was amplified by PCR and then cloned into the pGEX4T-2 (GST) to express the BnaA03.bZIP11 and BnaC03.bZIP11 protein in the *Escherichia coli* DE3 strain in the presence of 0.5 mM IPTG at 28 °C for 12 h. The recombinant BnaA03.bZIP11 and BnaC03.bZIP11 protein was purified using GST 4FF (Pre-Packed Gravity Column) (Sangon Biotech, C600911, Shanghai, China). For the electrophoretic mobility shift assay (EMSA), the Cy5-labeled probes and recombinant proteins were mixed in an EMSA/Gel-Shift binding buffer (Beyotime, GS005, China) at 25 °C for 20 min in the presence or absence of unlabeled competitor DNA. The reaction mixture was then electrophoresed on 6% non-denaturing polyacrylamide gels under ice-water conditions.

### Dual-luciferase assay

The full length of *Bna.bZIP11* was amplified by PCR and inserted into the *pGreenII-62SK* for transient overexpression. The 2-kb promoter sequence of *BnaA06.UF3GT* was also amplified by PCR and inserted into the *pGreenII 0800-LUC* vector using a ClonExpress II One Step Cloning Kit (Vazyme, C112, China). All relevant effector and reporter constructs were then transformed into Arabidopsis mesophyll protoplasts according to previously described methods [34]. The dual luciferase assay was then conducted according to the instructions from the Dual-Luciferase® Reporter Assay System (Promega, E1910, WI, USA). The data was expressed as the ratio of firefly to ranila luciferase activity (Fluc/Rluc). Each data point was based on at least three replicates, and three independent experiments were performed for each experiment.

### Functional enrichment analysis

Gene function descriptions were obtained from the *B. napus* (ZS11) reference genome. The GO enrichment analysis was performed using agriGO2, and terms with an FDR < 0.05 were considered significant.

### RT-qPCR assay

Quantitative reverse transcription PCR (RT-qPCR) assay was performed according to a previous report [35]. The melting curve of RT-qPCR was analyzed to ensure the presence of only one peak. The expression level of each gene was calculated using the 2^−ΔΔCT^ method. All analyses were performed at least three times. The *BnaActin7* gene (XM_013858992) was used as an internal control. All primers were listed in Table S1.

## Results

### Resynthesized allotriploid *B. napus* - *B. rapa* hybrids and phenotypic characterization

We previously generated two allotriploid hybrids of *Brassica* species by crossing two *B. napus* inbred lines (s*70*, A_s_A_s_C_s_C_s_; *yu25*, A_y_A_y_C_y_C_y_) with a *B. rapa* (Hort, A_h_A_h_) (see Materials and Methods). The two *B. napus* inbred lines had green stems, while the *B. rapa* had red stems (Fig. 1a). The resulting allotriploid hybrids, designated Hybrid-sh (*s70* × Hort, A_s_A_h_C_s_) and Hybrid-yh (*yu25* × Hort, A_y_A_h_C_y_), had 29 chromosomes and exhibited red stems similar to the *B. rapa* (Fig. 1a). Interestingly, Hybrid-sh had architectural features similar to *s70*, while Hybrid-yh showed characteristics between the two parentals (Fig. 1a). Additionally, the soluble protein and sugar contents in the two allotriploid F_1_ hybrids were approximately 141-168 mg/g and 51-56 mg/g higher, respectively, compared to their respective parental lines (Fig. S1a). The total phenolic content in the allotriploid hybrids was similar to that of the maternal lines and exceeded that of the paternal line (Student’s *t*-test, *p* < 0.05; Fig. S1a). The total flavonoid content in the allotriploid F_1_ hybrids was approximately 7.2-8.1 mg/g, which was higher than that of the maternal line but lower than that of the paternal line (Fig. S1b). Notably, the flavonoid content of Hybrid-yh was higher than that of Hybrid-sh (Student’s *t*-test, *p* < 0.05, Fig. S1b).

**Figure 1.**
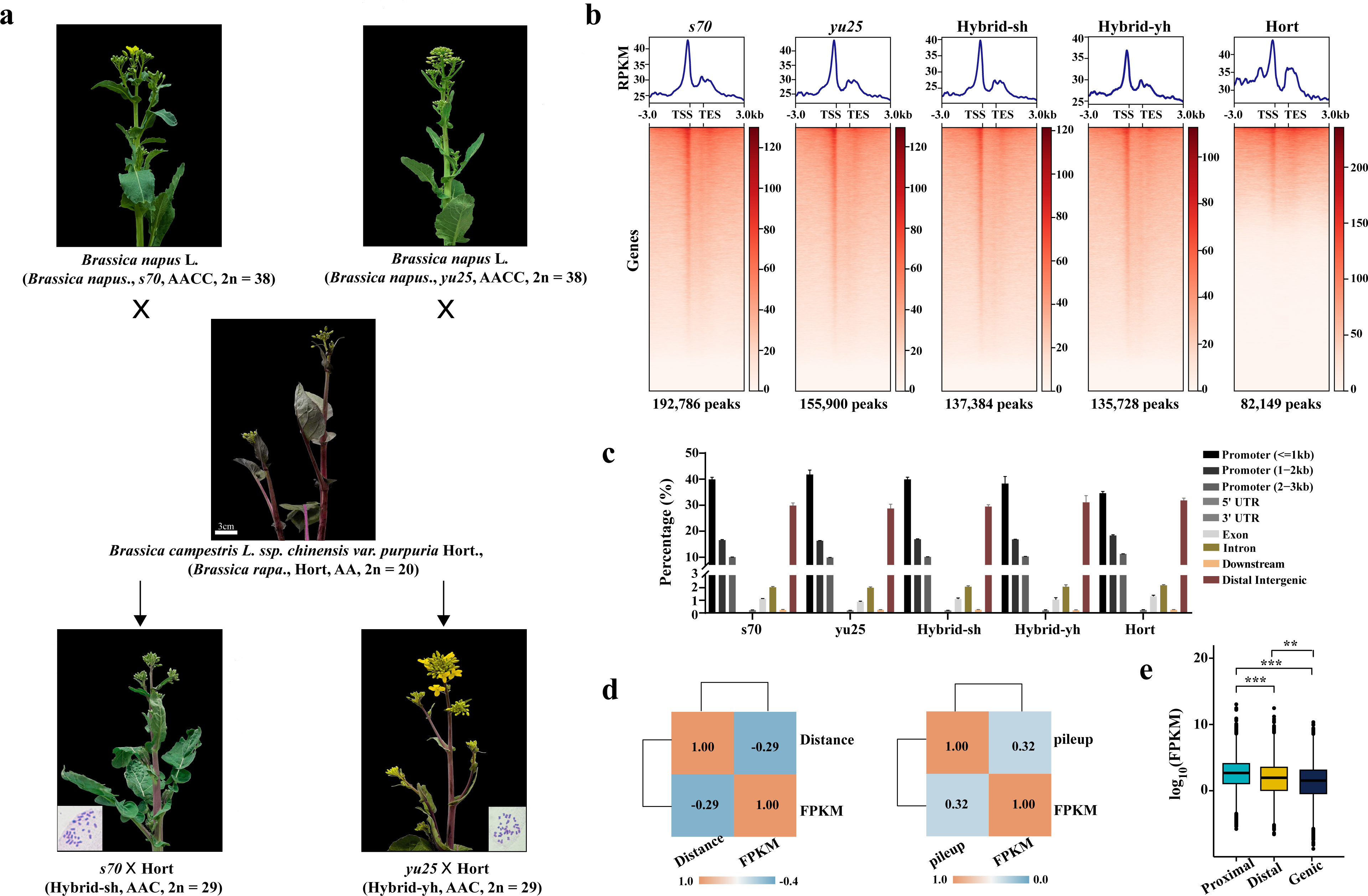
Model of hybridization-induced variation of accessible chromatin regions and DNA methylation sites in F_1_ hybrids. F_1_ hybrids inherit accessible chromatin regions (ACRs), sRNA, and DNA methylation loci from their parents, resulting in gene transgressive expression and single parental expression (SPE). Silent ACR: ACRs were identified in parents but not in F_1_ hybrids. Novel ACR: ACRs were identified in F_1_ hybrids but not in parents. Transgressive up genes: The gene expression level of the F_1_ hybrids was higher than that of the parents. Transgressive down genes: The gene expression level of F_1_ hybrids was lower than that of the parents. SPE: Genes are expressed only in one parent and F_1_ hybrids, but not in the other parent.

### Genome-wide identification of ACRs in F_1_ hybrids and their respective parentals

We then conducted RNA-seq and ATAC-seq of stem epidermis from two allotriploid hybrids (Hybrid-sh and Hybrid-yh) and their corresponding parental lines to investigate the genome-wide transcriptional dynamics during allotriploid hybridization. The RNA-seq generated an average of 20 million clean reads, with 93.4% successfully mapped to the reference genomes (Table S2). Meanwhile, ATAC-seq produced an average of 47 million clean reads per replicate (Table S3). Spearman’s rank correlation and principal component analysis (PCA) revealed high-quality RNA-seq and ATAC-seq datasets (Fig. S2a, S2b). A total of 192,786, 155,900, 137,384, 135,728, and 82,149 ATAC-seq peaks were enriched in *s70*, *yu25*, Hybrid-sh, Hybrid-yh, and Hort, respectively (Fig. 1b). The ACRs were significantly enriched at gene transcription start sites (TSS) (Fig. 1c). The majority of ACRs were between 200 and 500 base pairs (bp) in length, with the highest enrichment observed at 250 bp (Fig. S2c). Overall, the ATAC-seq data provide a comprehensive and reliable overview of ACRs in allotriploid hybrids and their parental plants.

The distribution of ACRs on the genome was categorized based on their proximity to genes, which were classified as Genic (overlapping with a gene), Proximal (TSS and TTS (transcription termination site), within 2 kb upstream or downstream of a gene, respectively), and Distal (more than 2 kb away from any gene). Our analysis revealed a positive correlation between gene expression levels and the presence of ACRs in the TSS region (*R*=0.32, Kruskal-Wallis test, *p* < 2.2e-16) (Fig. 1d). Conversely, gene expression showed a negative correlation with the distance of TSS region ACRs (*R*=-0.29, Kruskal-Wallis test, *p* < 2.2e-16) (Fig. 1d). On average, genes associated with Proximal (including TSS and TTS) ACRs were more highly expressed than the genes associated with Distal and Genic ACRs (Fig. 1e). In contrast to Distal and Genic ACRs, Proximal ACRs played a predominant role in regulating gene expression, suggesting that the positioning of ACRs is closely related to their transcriptional regulation, which is consistent with previous studies [36–38].

### ACRs show different convergent distributions after interspecific hybridization

Gene expression quantification analysis was then performed to examine the pre-existing levels of gene expression in the allotriploid hybrids and their parents. This involved comparing *in silico* hybrids (where the RNA-seq data combined maternal and paternal in a 1:1 ratio) with the F_1_ hybrids. The results showed that the majority of expressed genes exhibited additive expression patterns in both Hybrid-sh (82.9%) and Hybrid-yh (90.1%) (Fig. 2a). However, a total of 7,565 and 4,217 genes showed non-additive expression in Hybrid-sh and Hybrid-yh, respectively (Fig. 2a). The non-additively up-regulated genes in Hybrid-sh were mainly associated with responses to biotic and abiotic stimuli, immune responses, and metabolic processes (Table S4). Conversely, Hybrid-yh was enriched for ion transport, response to hormones, and metabolic processes (Table S5). Compared to their parents, the enrichment of process-related terms in the F_1_ hybrids suggests an enhanced capacity for metabolite production and improved tolerance to environmental conditions.

**Figure 2.**
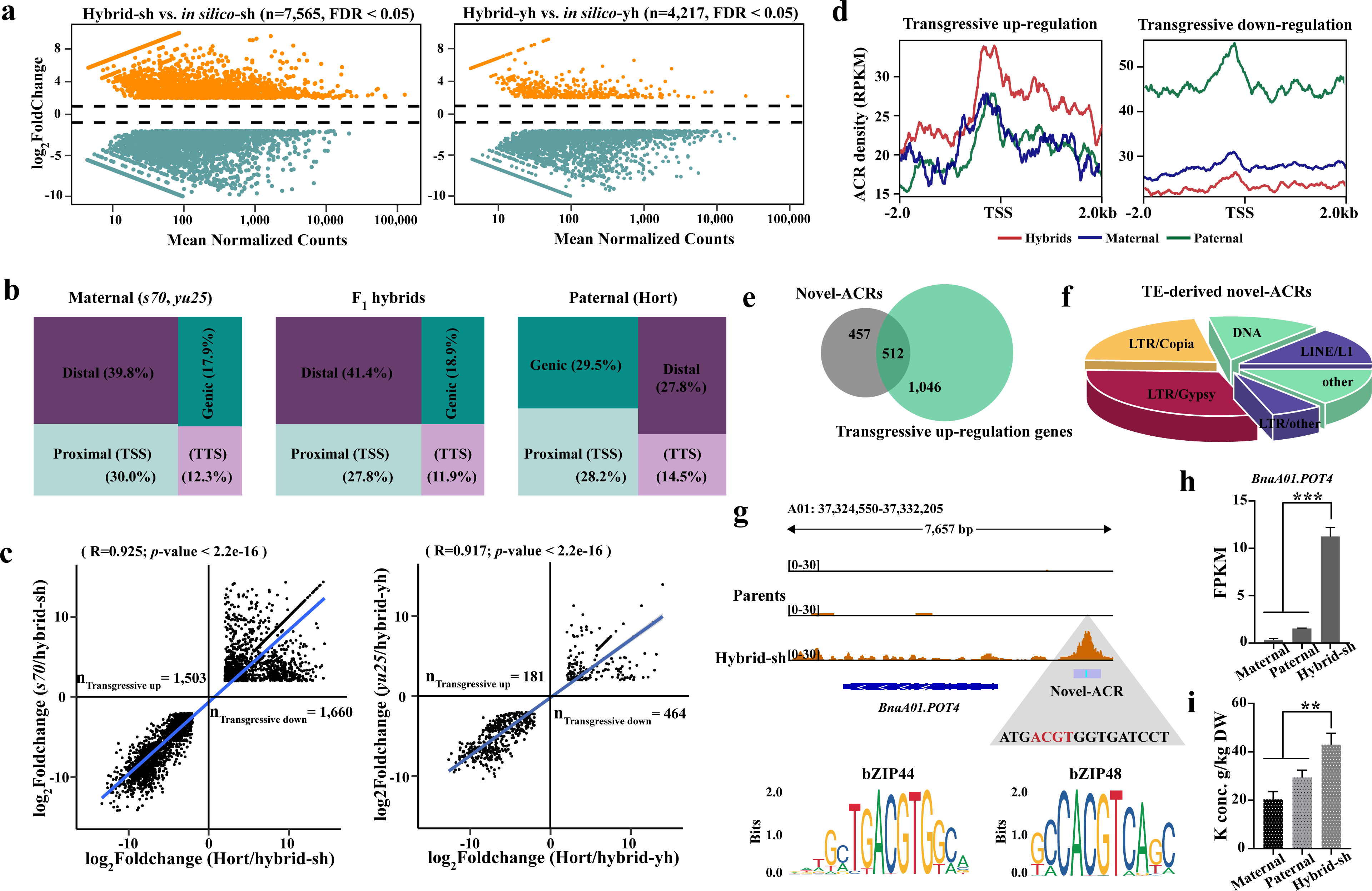
Overview of chromatin accessibility in the F_1_ hybrids and their parental. **(a)** The images showed the phenotypes of the allotriploid F_1_ hybrids and their parents at the flowering stage (scale bars=3 cm) and the typical chromosome number (Scale bars = 10 µm) of the allotriploid *Brassica* hybrids. **(b)** Chromatin accessibility around genes in the F_1_ hybrids (Hybrid-sh and Hybrid-yh), and their relative parents (maternal parents: *s70* and *yu25*; paternal parent: Hort). **(c)** The bar graph showed the proportion of ACR regions in the genome of F_1_ hybrids and their relative parents. **(d)** Comparison of gene expression and ACR intensity (left) and ACR distance (right). **(e)** Expression levels of genes associated with different types of ACRs. Boxplots showed the median (horizontal line). The upper and lower quartiles were the boundaries of the boxplots. The outlier was the data point outside the whiskers of the box plot. The Kruskal-Wallis test calculated the *p*-value (\*\**p* < 0.01; \*\*\**p* < 0.001).

After examining the variation in the distribution of ACRs within genes and across the genomes of parents and hybrids, it was found that more than 40% of the ACRs in the F_1_ hybrids were located in distal regions. In comparison, only 28% were distributed distally in the paternal line (Fig. 2b). After hybridization, there was a 2.8% decrease in Proximal ACRs, with 39.7% in F_1_ hybrids compared to 42.5% in the parents (Fig. S2d). Conversely, Distal ACRs showed a 6.7% increase in F_1_ hybrids compared to parental ACRs (Fig. S2e), confirming a distinct ACR distribution between parents and F_1_ hybrids. These results suggest that the hybridization process induces changes in the distribution and abundance of accessible chromatin regions, particularly in gene regions and their proximity.

### Transgressive gene expression mediated by novel ACRs

Previous studies have emphasized the potential role of non-additive genes in resynthesized plant materials [35,39]. In particular, transgressive genes represent an extreme form of non-additive genes, where their expression levels in the progeny exceed the highest or fall below the lowest levels observed in the parental lines [40]. Approximately 3,163 and 645 transgressive expression genes were identified in Hybrid-sh and Hybrid-yh, respectively (Fig. 2c). In Hybrid-yh, the number of down-regulated genes (464) was significantly higher than the number of up-regulated genes (181). In comparison, no difference was observed between the number of up-regulated and down-regulated transgressive genes in Hybrid-sh (Fig. 2c). Further investigation of these transgressive genes revealed 858 novel genes and 132 silent genes (Fig. S3a). These results suggest that the expression levels of genes in Hybrid-yh show relative stability in the hybrid offspring, with a greater likelihood of inheriting the expression pattern from the parent.

In F_1_ hybrids, chromatin accessibility was found to be higher than in the parents for transgressively up-regulated genes (Hybrids > Maternal = Paternal) (Fig. 2d). Conversely, for transgressive down-regulated genes, the chromatin accessibility was significantly lower in F_1_ hybrids than in the parents (Paternal > Maternal > Hybrids) (Fig. 2d). Further investigation revealed the presence of transgressive ACRs, including 969 novel ACRs and 25 silent ACRs (Fig. S3b), which were closely associated with the remodeling of gene expression regulation and the openness of chromatin accessibility. Notably, more than half of the novel ACRs were found to target transgressive up-regulated genes (Fig. 2e). Additionally, 375 (38.7%) of these novel ACRs were classified as TE-driven ACRs, with LTR/Gypsy-type retrotransposons playing a prominent role (30.4%) compared to other types of TEs (Fig. 2f). The presence of LTR and LINE elements significantly influences the evolution of novel ACRs in *Brassica* during hybridization. Furthermore, the chromosomal distribution of novel ACRs was mainly located at the telomeres, distal to the centromere (Fig. S3c).

The transgressive up-regulated genes in Hybrid-sh showed significant enrichment in metal ion binding, especially magnesium ion, calcium ion, and potassium ion binding, as well as secondary metabolic processes (Fig. S3d). In particular, two genes, *POT4* (*POTASSIUM TRANSPORTER 4*) and *POT2* (*POTASSIUM TRANSPORTER 2*), which are involved in the regulation of potassium ion uptake and transport in plants (Elumalai *et al*., 2002; Ahn *et al*., 2004), were identified. Two novel ACRs associated with *BnaA01.POT2* and *BnaA01.POT4* were identified in their promoters (Fig. 2g, Fig. S3e). The expression of *BnaA01.POT2* and *BnaA01.POT4* was significantly higher in Hybrid-sh compared to the parental lines (Fig. 2h, Fig. S3f). As expected, Hybrid-sh has a significantly higher potassium content than the parental lines (Fig. 2i). Previous studies have shown that the loss of the *bZIP48* gene function in rice results in increased sensitivity to zinc deficiency, while the *bZIP44* gene is involved in the plant’s response to cadmium tolerance [41,42]. Notably, the *BnaA01.POT4* novel ACR harbored *cis*-acting elements associated with the transcription factors *bZIP48* and *bZIP44*, which were linked to the ACGT *cis*-element (Fig. 2g). Based on these findings, we hypothesize that *bZIP48* and *bZIP44* may interact with *BnaA01.POT4* and thereby play a role in potassium ion transport.

To investigate the differences in mineral element content between two F_1_ hybrids and their parents, we analyzed 16 mineral elements, including five major elements (e.g., Mg) and eight trace elements (e.g., B) (Table S6). The results showed that the major elements (e.g., Ca) were significantly higher in F_1_ hybrids compared to their parents. However, the trace elements (e.g., B) showed additive effects, and other elements such as Fe, Zn, Cu, and Mo showed no significant differences between F_1_ hybrids and their parents. These results suggest that genetic factors regulate the mineral element content in F_1_ hybrids and are not solely determined by the additive effects of the parent plants.

### *Bna.bZIP11* in SPA-ACR regulates the expression of *BnaA06.UF3GT* to promote the accumulation of anthocyanins

The *c*-means clustering method was then used to classify the differential peaks according to chromatin accessibility levels in F_1_ hybrids and their respective parents, resulting in nine clusters labeled C1 to C9 (Fig. 3a, Fig. S4a). For ACRs, the ATAC-seq peaks were detected in only one parent, while they were not detected in the other parent, termed single-parent activation ACRs (SPA-ACRs). These SPA-ACRs could be further categorized into two patterns: SPA-M ACRs, where the ACRs were detected in the maternal and F_1_ hybrids but not in the paternal, and SPA-P ACRs, where the ACRs were detected in the paternal and F_1_ hybrids but not in the maternal. Since the Hort lacks the C-subgenome, it’s reasonable to expect a limited number of ATAC-seq peaks in the Hort materials compared to those in the maternal (AACC) and F_1_ hybrid (AAC) genomes. We then focused the analysis only on peaks corresponding to the A-subgenomes. A total of 8,125, 9,681, and 22,885 SPA-ACRs were identified in the A_s_ (A-subgenome of *s70*, *B. napus*), A_y_ (A-subgenome of *yu25*, *B. napus*), and A_h_ (A genome of *B. rapa*) subgenomes (Fig. 3a, Fig. S4a), which were represented in cluster C5 and cluster C8, respectively (Fig. 3a, Fig. S4a). The SPA-ACR can lead to the emergence of single parental expression (SPE), a gene is expressed in only one parent, while it is silent in the other parent after interspecific hybridization [34]. On average, 336 genes expressed in the F_1_ hybrids were exclusively expressed in the maternal parents (SPE-M, A_s_ and A_y_), and 277 genes expressed in the F_1_ hybrids were exclusively expressed in the paternal parent (SPE-P) in the A genome (A_h_) (Fig. S4b). Approximately, 20% of the SPE genes were correlated with SPA-ACR (Fig. S4c).

**Figure 3.**
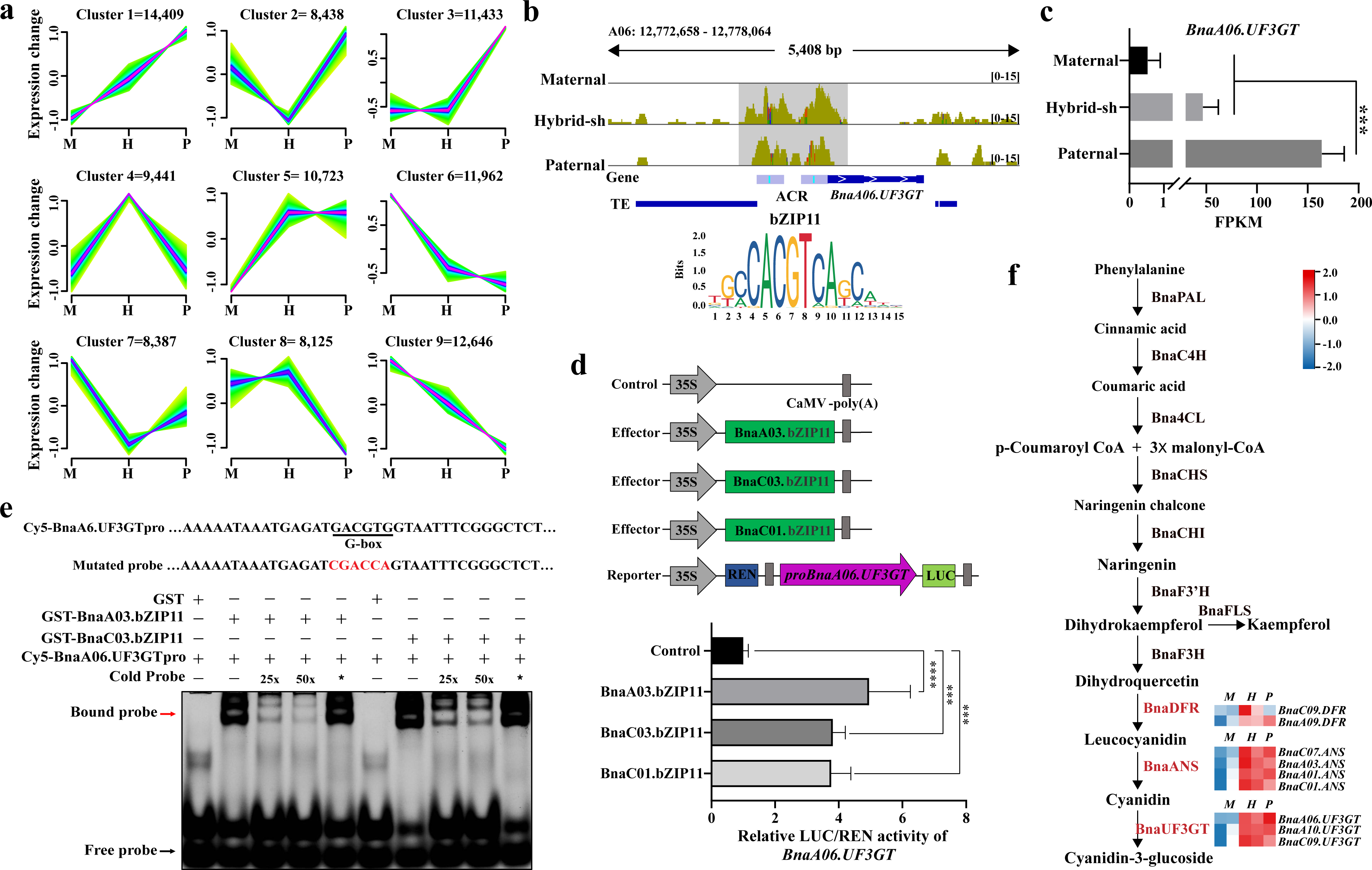
Chromatin accessibility and the expression of transgressive genes in the F_1_ hybirds. **(a)** The MA-plot showed differentially expressed genes (DEGs) between the F_1_ hybrids and the mid-parent value (MPV). **(b)** The graph showed the proportion of ACR regions in Genic (overlapping with a gene), Proximal [within 2 kb upstream or downstream of a gene, including transcription start site (TSS) and transcription termination site (TTS)], and Distal (more than 2 kb away from any gene) in the F_1_ hybrids and their parents. **(c)** The scatterplot showed the distribution of differential expression levels of Hybrid-sh (left) and Hybrid-yh (right) versus maternal (*x*-axis) and paternal (*y*-axis) parents. **(d)** The graph showed the ACR densities of transgressive up-regulated (left) and down-regulated (right) genes. **(e)** Venn diagram showed an overlap of novel ACRs and transgressive up-regulated genes. **(f)** The pie chart showed the proportion of TE-derived novel ACRs (TE-driven ACRs when more than 50% of the region overlaps with TEs). **(g)** The genome browser showed ATAC-seq peaks around *Potassium Transporter 4* (*POT4*) in the parents and Hybrid-sh, and the enrichment of the *bZIP44* and *bZIP48* motif (ACGT) in the promoter region of *POT4*. **(h)** and **(i)**, the bar graph showed the expression level of *BnaA01.POT4* **(h)**, and potassium levels in Hybrid-sh and its relative parents. Error bars indicated the mean ± SD of three biological replicates. In **(h)** and **(i)**, the Student’s t-test was used to calculate significance where ** and *** indicated *p* < 0.01 and *p* < 0.001, respectively.

The datasets were then prioritized based on the occurrence of Proximal SPA-P ACRs and SPE-P genes. Notably, the top 10 ranked in the SPA-P genes showed a significant presence of genes associated with anthocyanin synthesis (Fig. S4d). For example, we found that the chromatin accessibility of *BnaA06.UF3GT*, which encodes the flavonoid 3-O-glucosyltransferase (UF3GT) enzyme [43], was more significant in the hybrids and the paternal promoter region compared to the maternal promoter region (Fig. 3b). The expression of the *BnaA06.UF3GT* was found to be significant in F_1_ hybrids and paternal individuals, while no expression was detected in the maternal plant (Fig. 3c). Analysis using the HOMER algorithm showed that the *BnaA06.UF3GT* gene might be regulated by the *BnabZIP11* transcription factor (Fig. 3b). To investigate this, a luciferase (LUC) gene driven by a ∼2 kb promoter of *BnaA06.UF3GT* was used as a reporter, and three *BnabZIP11* genes were driven by the *CaMV 35S* promoter (Fig. 3d). Co-transformation of the vector containing *BnaA03.bZIP11*, *BnaC03.bZIP11*, and *BnaC01.bZIP11* with the *pBnaA06.UF3TG-LUC* construct significantly increased LUC/REN activities by 4.95, 3.81, and 3.76-fold, respectively, compared to the expression of *pBnaA06.UF3TG-LUC* alone (Fig. 3d). Further analysis revealed the binding of the BnabZIP11 transcription factor to the G-box elements upstream of *BnaA06.UF3GT* (Fig. 3e), suggesting that *BnabZIP11* may directly bind to the G-box elements upstream of the *BnaA06.UF3GT* promoter and thereby positively regulate the expression of *BnaA06.UF3TG*. Additionally, the expression patterns of genes involved in anthocyanin synthesis and related transcription factors were also highly expressed in the F_1_ hybrids and the paternal parent (Fig. 3f, Fig. S4e). This finding suggests a possible reason for the high accumulation of total anthocyanins and phenolics in the paternal and hybrid plants (Fig. S1b). The activation of ACRs by a single parent after interspecific hybrid genome recombination is positively associated with gene regulation, which may contribute to the non-additive phenotypic traits observed in resynthesized F_1_ hybrids. The interaction between the activated ACRs and gene expression patterns may lead to altered gene regulatory networks, resulting in unique phenotypic traits in hybrids that differ from those of either parent.

### Hybridization-induced DNA methylation related to parents in F_1_ hybrids

The process of hybridization often results in significant reprogramming of global DNA methylation. This can be considered an “epigenetic shock” resulting from the fusion of different epigenomes from the parental plants [44,45]. Therefore, we evaluated the total methylation levels in hybrids and compared them with the parental methylation levels. The sequencing depth of WGBS was approximately 30-fold genome coverage, and the bisulfite conversion was greater than 99% in all samples (Table S7). The sRNA-seq data generated an average of 20 million clean reads per replicate, of which 93.4% were mapped to the reference genome (Table S8). PCA plots and Spearman’s rank correlation coefficients generated from the sRNA-seq and WGBS libraries showed clear clustering patterns and reproducibility across biological replicates (Fig. S5a, Fig. S5b). At the chromosomal level, a bias in the total methylation levels of CG, CHG, and CHH towards the hypermethylated parent was observed (Fig. S6a). This indicates that the hybrid plants tended to inherit higher methylation levels from the parent with higher methylation at these specific sequence contexts.

To investigate the changes in DNA methylation in F_1_ hybrids after interspecific hybridization, two *in silico* hybrids (*in silico*-sh and *in silico*-yh) were first constructed by combining maternal and paternal WGBS data in a 1:1 ratio. All DNA methylation levels (CG, CHG, and CHH) of Hybrid-sh were higher than those of *in silico*-sh, while no significant difference in the methylation levels of CG and CHG was found between Hybrid-yh and *in silico*-yh (Fig. 4a). However, the methylation level of CHH was lower in Hybrid-yh than *in silico*-yh (Fig. 4a). Compared to *in silico* hybrids, 24,693 and 9,214 DMRs (Differentially Methylated Regions) were identified in Hybrid-sh and Hybrid-yh, respectively (Fig. 4b). There were fewer hypo-DMRs (9,701) compared to hyper-DMRs (13,492) in Hybrid-sh (Fig. S6b). 9,214 DMRs were found in Hybrid-yh, and nearly half of the DMRs (49%) were CHH-DMRs (Fig. S6b). Compared to Proximal and Distal DMRs, a greater number of Genic DMRs were identified for CG methylation in both Hybrid-sh and Hybrid-yh (Fig. 4b). For CHG methylation, the distribution of DMRs was as follows: Distal > Proximal > Genic (Fig. 4b). The distribution of DMRs at gene locations suggests that this distribution is influenced by the context of DNA methylation. More than 68% of the differentially methylated genes (DMGs) were Proximal DMGs (Fig. 4c). Transcript levels of Proximal DMGs were then compared between F_1_ hybrids and *in silico* hybrids. No significant difference was observed in the expression of DMGs associated with CHG (Fig. 4d). However, the expression of hyper DMGs of CHH was significantly higher in F_1_ hybrids compared to *in silico* hybrids, while hypo DMGs showed the opposite trend (Fig. 4d).

**Figure 4.**
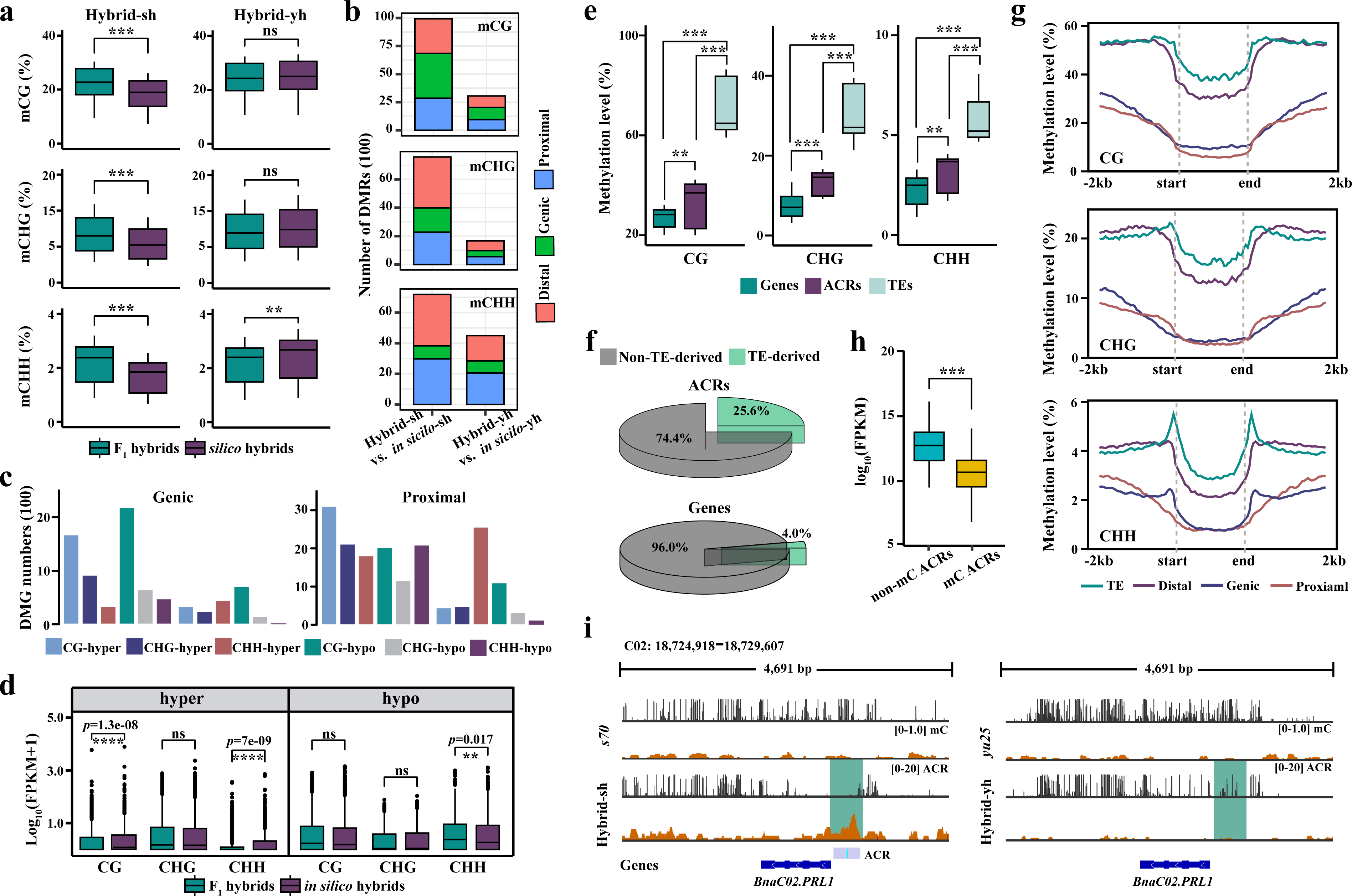
A-subgenome SPA-ACR drives bZIP11 to promote UF3GT expression in paternal and Hybrid-sh. **(a)** The graphs showed the *c*-means soft clustering analysis of chromatin accessibility levels of A-subgenomes in the Hybrid-sh and its relative parents. **(b)** The genome browser showed the ATAC-seq peaks around *BnaA06.UF3GT* in the parents and Hybrid-sh, and the bZIP11 motif (ACGT) enrichment in the promoter region of *BnaA06.UF3GT*. **(c)** The bar graph showed the expression level of *BnaA06.UF3GT* in the Hybrid-sh and its relative parents. **(d)** Schematic diagrams of the effector and reporter constructs for the dual-luciferase transcriptional activity assay. The bar graph showed the LUC/REN activities of Arabidopsis protoplasts after co-transformation with the (p*BnaA06.UF3GT:LUC*) and reporter constructs (p*35S* (control), and p*35S:BnaA03.bZIP11* (*BnaC03.bZIP11* and *BnaC01.bZIP11*)). **(e)** The EMSA results showed that BnaA03.bZIP11 and BnaC03.bZIP11 could directly bind to the *BnaA06.UF3GT* promoter. Increasing amounts (25- and 50-fold) of the unlabeled DNA fragments were added as competitors. The red arrow indicated the shift bands. Red letters indicate the mutated G-box cis-element within the *BnaA06.UF3GT* promoter. An asterisk indicates that the mutated cold competitor probes were added to the panel. **(f)** Anthocyanin biosynthesis pathway. The colored boxes were the gene expression heatmap from RNA-seq, normalized by log_2_FPKM (calculation method). Genes responsible for Leucocyanidin, Cyanidin, and Cyanidin-3-glucoside steps were shown in red. BnaPAL, phenylalanine ammonialyase; BnaC4H, cinnamic acid 4-hydroxylase; Bna4CL, 4-coumarate:coenzyme A ligase; BnaCHS, chalcone synthase; BnaCHI, chalcone isomerase; BnaF3H, flavanone 3-hydroxylase; BnaF3ʹH, flavonoid 3ʹ-hydroxylase; BnaDFR, dihydroflavonol-4-reductase; BnaANS, anthocyanidin synthase; BnaUF3GT, UDP glucose-flavonoid 3-O-glucosyltransferase. The colored boxes are the heatmap of gene expression from RNA-seq normalized by log_2_FPKM. In **(h)** and **(i)**, error bars indicated the mean ± SD of three biological replicates. Student’s t-test was used to calculate significance, with *** indicating *p* < 0.001.

TE involvement was detected in approximately 55.6% of DMRs, further analysis revealed that 21.5% and 39.1% of these DMRs were LINE and LTR (Fig. S6c). To gain further insight into the relationship between sRNA-mediated DNA methylation and hybridization-induced hyper- and hypo-DMRs, we analyzed changes in sRNA enrichment within hyper- and hypo-DMRs. In hyper-DMRs, we observed a significant increase in sRNA accumulation *in silico* compared to hybrids, whereas the opposite trend was seen in hypo-DMRs (Fig. S6d). This highlights the close relationship between DMRs and sRNA accumulation.

### DNA methylation in ACR associated with TEs in F_1_ hybrids

Previous reports indicate that DNA methylation plays a critical role in the maintenance and loss of chromatin accessibility [13]. When comparing the DNA methylation levels of TEs, gene bodies, and ACRs, we found that ACRs had significantly higher methylation levels compared to gene bodies, but lower than TEs (Kruskal-Wallis test, *p* < 2.2e-16, Fig. 4e). Further analysis revealed that 25.6% of ACRs and 4.0% of genes were derived from TEs (Fig. 4f), indicating a substantial overlap between ACRs and TEs. This finding suggests that the increased DNA methylation levels observed in ACRs may be associated with TEs, potentially explaining the observed differences between ACRs and gene bodies.

The level of DNA methylation in ACRs varies depending on their position in the genome. ACRs with a CG context show different levels of DNA methylation, with a consistent decrease in methylation levels between TEs, Distal, Genic, and Proximal regions (TE > Distal > Genic > Proximal; Kruskal−Wallis, *p* < 2.2e−16; Fig. 4g). However, in the context of CHG and CHH, there was no significant difference between the methylation levels of the Genic and Proximal regions (Fig. 4g). Overall, TE and Distal ACR showed significantly higher methylation levels compared to Genic and Proximal ACRs (Fig. 4g). This suggests that TE and distal regions may require higher levels of DNA methylation for stability. In contrast, genic and proximal regions may be more sensitive to gene expression regulation, resulting in lower levels of DNA methylation. Gene expression analysis also supported this observation, as the expression of genes associated with ACRs lacking DNA methylation was significantly higher compared to genes associated with methylated ACRs (Fig. 4h). For example, in Hybrid-sh, an open chromatin region was observed in the promoter region of *BnaC02.PRL1*, which did not inherit the methylation sites from the parent; conversely, in Hybrid-yh, the situation was reversed (Fig. 4i). RNA-seq and RT-qPCR results showed that *BnaC02.PRL1* was not detected in Hybrid-yh, but was significantly expressed in Hybrid-sh (Student’s *t*-test, *p* < 0.05, Fig. S6e). This indicates that DNA methylation plays a role in repressing gene expression within ACRs, and unmethylated ACRs may be more transcriptionally active.

### Genome dosage affects accessible chromatin regions in F_1_ hybrids

Genome-wide dose-dependent and independent regulation contributes to the evolution and gene expression of plant polyploids [46], and genome imbalance in AAC hybrids may lead to changes in gene expression and variation in epigenetics. We then calculated the correlation coefficient (*R*-values) between the expression of 96,992 genes and the relative doses of the genes in the A and C-subgenomes. 19,501-25,011 and 22,554-26,420 dose-dependent genes were identified in the A and C-subgenomes, respectively (Fig. 5a). Interestingly, approximately 56% and 16% of the dose-dependent and dose-independent genes, respectively, were consistent in the two F_1_ hybrids (Fig. S7a), indicating that dose-dependent genes are more conserved in the F_1_ hybrids.

**Figure 5.**
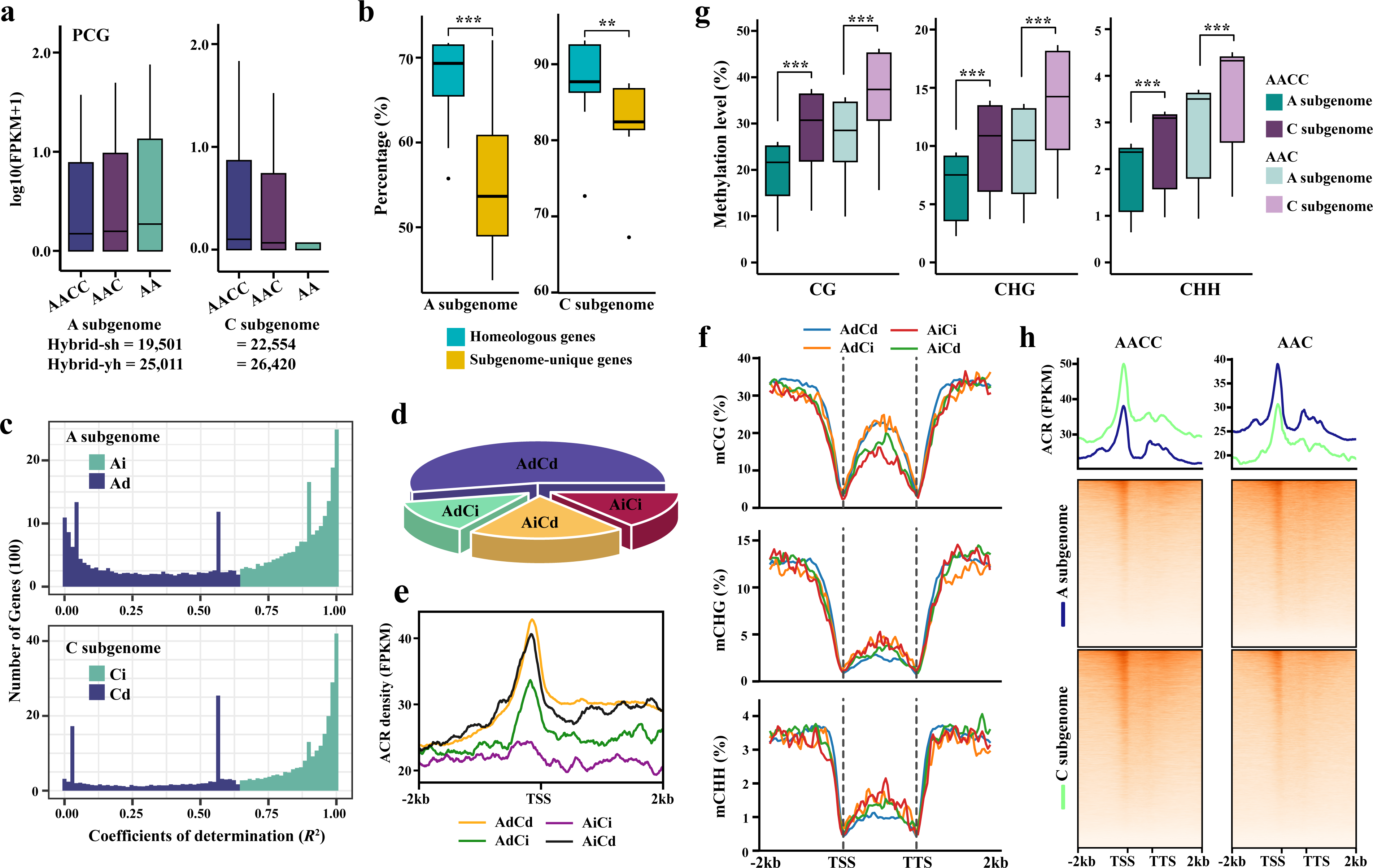
Divergence of DNA methylation landscape between parents and hybrids. **(a)** The boxplots showed the total DNA methylation level (CG, CHG, and CHH) of F_1_ hybrids (Hybird-sh (left) and Hybird-yh (right)) and *in silico* hybrids (Wilcoxon rank-sum test; \*\**p* < 0.01; \*\*\**p* < 0.001; ns, no significant difference). **(b)** The bar graph showed the number of DMRs between F_1_ hybrids and *in silico* hybrids of 200 bp bins. Different colors indicate the distribution of DMRs in the distal (red), genic (green), and proximal (blue) regions. **(c)** The bar graph showed the number of Genic (left) and Proximal (right) DMGs in the Hybrid-sh and Hybird-yh. **(d)** Boxplots showed the expression levels of hyper- and hypo-DMGs in F_1_ hybrids and *in silico* hybrids with methylation sites in proximal regions. The *y*-axis represented the gene expression level log_10_(FPKM+1) (Wilcoxon rank-sum test; \*\**p* < 0.01; \*\*\**p* < 0.001; ns, no significant difference). **(e)** The boxplots showed the methylation level (CG, CHG, and CHH) across genes, TEs, and ACRs. **(f)** The pie chart showed the proportion of TE-derived and non-TE-derived ACRs (top) and genes (bottom). The TE-derived ACR was defined as having more than 50% of the region overlapping with TEs. **(g)** The image showed the methylation level (CG, CHG, and CHH) of TE ACRs, Distal ACRs, Genic ACRs, and Proximal ACRs. **(h)** The boxplots showed the expression level of genes that associated with methylated and non-methylated ACRs. **(i)** The genome browser showed DNA methylation loci around *BnaC02.PRL1* in F_1_ hybrids (Hybrid-sh and Hybrid-yh) and their relative parents. In **(e)** and **(h)**, The upper and lower quartiles were boundaries of the boxplots. The Kruskal-Wallis test calculated the *p*-value (\*\**p* < 0.01; \*\*\**p* < 0.001).

**Figure 6.**
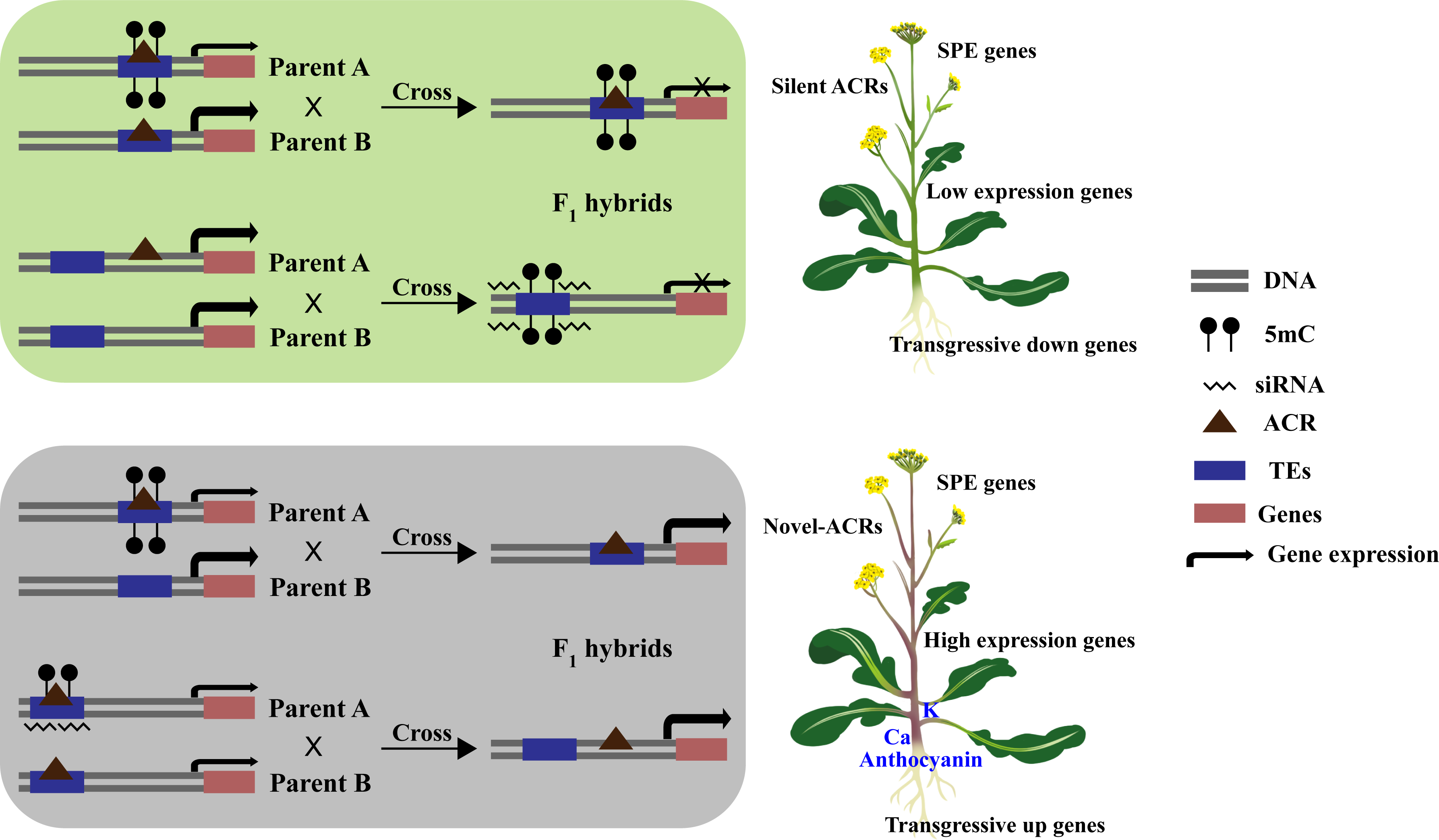
Effects of genomic imbalance on accessible chromatin regions and DNA methylation in F_1_ hybrids. **(a)** The expression level (top) and number (bottom) of dose-dependent genes in the A and C-subgenomes. **(b)** The boxplots showed the proportion of dose-dependent homologous gene pairs and dose-dependent genome-unique genes in the A and C-subgenomes. **(c)** The graph showed the number of dose-dependent and dose-independent homologous genes in the A and C-subgenomes. Pearson’s correlation was used to test and divide all homologous genes into two groups based on the coefficient of determination (R^2^) to compare the characteristics of dose-dependent and dose-independent homologous genes. Homologous genes with statistically significant correlations were designated as dose-dependent A (Ad) and C (Cd) genes, while those with insignificant correlations were designated as dose-independent A (Ai) and C (Ci) genes. **(d)** The pie chart showed the percentage of AdCd, AdCi, AiCd, and AiCi duplicated genes. **(e)** and **(f)**, the images showed the ACR density **(e)** and DNA methylation level (CG, CHG, and CHH) **(f)** between AdCd, AdCi, AiCd, and AiCi duplicated genes. **(g)** and **(h)**, the images showed the distribution of DNA methylation level **(g)** and ACR density **(h)** of A and C-subgenomes in AACC and AAC.

Pearson’s correlation test was used to divide all homologous genes into two groups based on the coefficient of determination (*R*^2^) to compare the characteristics of dose-dependent and dose-independent homologous genes. Homologous genes with statistically significant correlations were designated as dose-dependent A (Ad) and C (Cd) genes, while those with insignificant correlations were designated as dose-independent A (Ai) and C (Ci) genes. The proportion of dose-dependent genes among the homologous genes in the A/C-subgenome was higher than that of genome-unique genes (Fig. 5b). Among the four categories of homologous genes (AdCd, AdCi, AiCd, and AiCi), AdCd had the highest number (20,888), followed by AiCd (7,746), while AdCi and AiCi had the lowest number (4,927 and 5,552, respectively) (Fig. 5c, Fig. 5d). Chromatin stacking was observed to be highest in the ACR for AdCd, followed by AdCi, and lowest for AiCi (Fig. 5e), indicating that dose-dependent homologous genes have increased chromatin accessibility, making them more susceptible to regulation by transcriptional and regulatory factors. Higher levels of DNA methylation were found in the CG context for AdCd and AdCi compared to AiCd and AiCi, while in the CHG and CHH contexts, AdCd had the lowest methylation levels among the four categories and AiCi had the highest (Fig. 5f). This suggests that different gene categories may be regulated by different DNA methylation modifications. In both AACC and AAC, the methylation level of the C-subgenome was higher than that of the A-subgenome, regardless of whether the dose of the C-subgenome (Fig. 5g). This suggests that DNA methylation levels are regulated by factors beyond genomic dosage. In AACC, the ACR level of the C-subgenome was higher than that of the A-subgenome; however, in AAC, the ACR level of the A-subgenome was significantly higher than that of the C-subgenome (Fig. 5h). This demonstrates that in AAC, doubling the dosage of the A-subgenome results in a significant increase in the ACR level of the A-subgenome, highlighting the strong effect of genome dosage on accessible chromatin regions. Dose-dependent and dose-independent genes were identified in the A- and C-subgenomes. The dose-dependent genes exhibited higher chromatin openness. Notably, differences in the A-subgenome were specifically located in the TSS and upstream regions, whereas differences in the C-subgenome were observed more broadly across the genome (Fig. S7b).

## Discussion

Interspecific hybridization is essential in plant breeding and genetic improvement [7,47]. In this study, we resynthesized two interspecific F_1_ hybrids by crossing different genotypes of *B. napus* and *B. rapa*, to elucidate the intricate relationship between accessible chromatin regions, DNA methylation, and the expression of transgressive genes. The interplay between DNA methylation, TEs, and sRNA contributes to the dynamic landscape of ACRs during interspecific hybridization, resulting in distinct gene expression patterns across the genome.

### Accessible chromatin regions and DNA methylation differ in genome dosage effects

Chromatin accessibility is essential for gene transcription as it allows transcription factors and RNA polymerase to bind to DNA and initiate gene expression [48]. Changes in chromatin accessibility can directly impact gene expression levels, leading to significant dosage effects [37]. In contrast, DNA methylation has a more indirect effect on gene expression. It can influence gene expression through mechanisms like inhibiting transcription factor binding and recruiting inhibitory protein complexes [49–51]. These mechanisms may interact complexly and involve feedback regulation, making the relationship between DNA methylation and gene expression non-linear and resulting in an insignificant dose effect. Furthermore, DNA methylation patterns are dynamic and highly specific to different cell types [52]. Despite alterations in genome dosage, DNA methylation levels can remain relatively stable or change only in certain cell types. On the other hand, chromatin accessibility is also dynamic and cell-type specific, but it is more responsive to changes in genome dosage [53,54]. This is due to the fact that alterations in chromatin structure serve as a rapid mechanism for regulating gene expression and can more directly respond to changes in genome dosage.

### Interspecific hybridization triggers genome recombination leading to novel accessible chromatin regions

During plant hybridization, the unequal retention of epigenetic marks, particularly accessibility chromatin regions (ACRs) activated by single parents or novel, can significantly affect gene expression and the trait development in hybrid offspring. In *Camellia sinensis*, a large number of accessible chromatin regions is observed after interspecific hybridization [10]. The ‘newborn DHS’ (nbDHS) after cotton polyploidization suggests that these nbDHS may arise from transposable elements during or after polyploidization [36]. We found 512 novel ACRs near the transgressive genes in F_1_ hybrids, 38.7% of these novel ACRs were classified as TE-driven ACRs, suggesting that interspecific hybridization triggers genome recombination that activates ACR formation. SPA-ACRs suggest that only one parent may control the expression pattern of certain gene regions during genome recombination. The maintenance of this expression pattern may be influenced by asymmetric chromatin openness, which can lead to increased expression or silencing of specific genes in hybrid offspring [55]. Notably, most SPA-ACRs are found within the gene bodies that encode the proteins, suggesting a possible relationship between the expression pattern of single-parent activation and structural features of the gene’s transcriptional activity region. In addition, the higher expression levels of genes adjacent to SPA-ACRs provide further evidence of a direct link between this expression pattern and increased gene expression. In wheat, expression of the *ph1* gene is associated with chromatin remodeling, which alters chromatin structure so that only genes from a particular parent are expressed [56]. Differences in the proportions of DNase I hypersensitive sites (DHS) in distal and proximal regions have been observed during wheat polyploidization, suggesting that the distribution of DHS varies among different genomes [9]. Our results indicate that interspecific hybridization induces changes in the distribution and abundance of chromatin regions accessible to genic and proximal regions.

### Accessible chromatin regions and DNA methylation jointly regulate gene expression

There is a close relationship between DNA methylation and open chromatin regions [13,57]. DNA methylation typically occurs on CpG islands, where its primary role is to suppress gene expression. Methylated CpG islands often result in a more condensed chromatin structure, limiting the binding of transcription factors and other regulatory proteins and ultimately repressing gene expression [49,58]. In contrast, open chromatin regions have a more relaxed chromatin structure, allowing easier binding of transcription factors and regulatory proteins, thus promoting gene expression. Significant positive correlations have been observed between chromatin accessibility and gene expression levels in nearby genes in various plant species, such as sorghum, rice, and Arabidopsis [37,38,54]. We found that the DNA methylation level of the ACR located at the proximal end of the gene is much lower than that of the TE and distal regions (Fig. 4g). This may be because the proximal ACR usually contains promoters and enhancers and is an important region for gene transcription regulation. Low levels of DNA methylation help keep these regions open, making it easier for transcription factors and other regulatory proteins to bind, thereby promoting gene expression [16,57]. Variations in DNA methylation found in lettuce may affect chromatin accessibility, thereby altering the expression levels of associated proximal and distal genes [16]. While DNA methylation leads to chromatin compaction and gene inhibition by recruiting inhibitory protein complexes, the open chromatin structure allows for the action of DNA demethylases, leading to decreased DNA methylation levels [13,49]. This delicate balance is essential for maintaining normal gene expression patterns.

## Conclusions

Overall, our findings provide valuable insights into the role of accessible chromatin and DNA methylation in driving gene expression in hybrids (Fig. 1). We observed significant differences in the expression of non-additive genes among the different F_1_ hybrids, with up-regulated transgressive genes associated with metal ion accumulation and SPE genes associated with anthocyanin accumulation. This study contributes to our preliminary understanding of how accessible chromatin regions and DNA methylation regulate metabolite and ion accumulation in F_1_ hybrids.

## Ethical statement

This article does not contain any studies with human or animal subjects.

## Acknowledgments

We thank the National Key Laboratory of Crop Genetic Improvement of Huazhong Agricultural University for providing the bioinformatics computing platform, Dr. Shengwei Dou from Shandong Agricultural University for the experimental support, and Novogene provides sequencing services. This work was supported by the Science and Technology Innovation 2030-Major Project (2023ZD04068) to Cheng Dai and the National Natural Science Foundation of China (No. 32172070) to Chaozhi Ma.

## Author contributions

**Cheng Dai, Chengtao Quan, and Chaozhi Ma:** conceived the original idea. **Qin Zhang, Xiaoni Zhang, and Kexin Chai:** performed experiments; **Chengtao Quan and Guoting Cheng:** designed the work and analyzed the data; **Chengtao Quan and Cheng Dai:** wrote and discussed the paper with all authors.

## Data availability

All sequence data generated in this study have been deposited in the BIG data under the BioProject accession number PRJCA023096.

## Supporting Information

### Supplemental materials and methods

The details of ATAC-seq, WGBS, RNA-seq, sRNA-seq experiment, and statistics methods were described in supplemental materials.

