## Supplemental table for "Accessible chromatin regions and DNA methylation regulate gene expression leading to changes in agronomic traits in *Brassica* allotriploid hybrids"

**Title Page**

**Interspecific hybridization in Brassica species leads to changes in agronomic traits through the regulation of gene expression by chromatin accessibility and DNA methylation**

Chengtao Quan^1,2^, Qin Zhang^1,2^, Xiaoni Zhang^1,2^, Kexin Chai^1,2^, Guoting Cheng^3^, Chaozhi Ma^1,2^, and Cheng Dai^1,2^

1 National Key Laboratory of Crop Genetic Improvement, Huazhong Agricultural University, Wuhan 430070, China

2 Hubei Hongshan Laboratory, Wuhan, 430070, China

3 College of Informatics, Huazhong Agricultural University, Wuhan 430070, China.

The author responsible for the distribution of materials integral to the findings presented in this article following the policy described in the Instructions for Authors is:

Cheng Dai

To whom correspondence should be addressed.

Dr. Cheng Dai

National Key Laboratory of Crop Genetic Improvement, Huazhong Agricultural University, Wuhan 430070, P.R. China

**Supplemental materials and methods**

**ATAC-seq**

For each biological replicate, the collected plant tissue was cut into small pieces with blade in 500 mL lysis buffer (15 mM Tris-HCl pH7.5, 20 mM NaCl, 80 mM KCl, 0.5 mM spermidine, 5 mM 2-mercaptoethanol and 0.2% Triton X-100). After confirming nuclear integrity, purified nuclei were resuspended in a 50 μL Tn5 transposase integration reaction and incubated at 37°C for 30 min. The Tn5 transposase-digested DNA fragments were then recovered using a MinElute PCR Purification Kit (Qiagen, Cat No./ID: 28004), and followed by purification and amplification. The purified library was then sequenced on an Illumina Novaseq platform by Novogene Gene Technology (Novogene, Beijing, China). All ATAC-seq profiles were generated from at least of three independent biological replicates.

**Whole genome bisulfite sequencing**

Genomic DNA was extracted from 10 samples using the cetyl trimethylammonium bromide (CTAB) method. A lambda DNA spike-in was utilized to correct for non-conversion rates of uracil, with 1 ng of methyl-free lambda DNA added to 1 μg genomic DNA as an internal reference for the conversion test. Bisulfite conversion of DNA was carried out using the EZ DNA Methylation Gold Kit (Zymo Research in Irvine, California, USA). The Bisulfite-Seq Library Prep Kit for Illumina (Novogene in Beijing, China) was used to construct whole genome bisulfite sequencing (WGBS) libraries, which were then sequenced on an Illumina HiSeq X10 platform at a depth of 30-fold. Two biological replicates were performed.

**RNA-seq and sRNA-seq**

Total RNA was extracted with the RNeasy Plant Mini Kit (Qiagen, Cat No./ID: 74904) according to the manufacturer's instructions. This preparation was split and used for both RNA-seq and sRNA-seq. Library construction and deep sequencing were performed using the Illumina HiSeq 4000 Platform according to the manufacturer’s protocols (Novogene, Beijing, China). Two or three biological replicates were performed.

**Statistics and reproducibility**

Statistical significance was determined using R (https://r-project.org). The Wilcoxon rank sum test and the χ2 test were performed using the *Wilcoxon.test* function and the *chisq.test* function, respectively, from the R package.

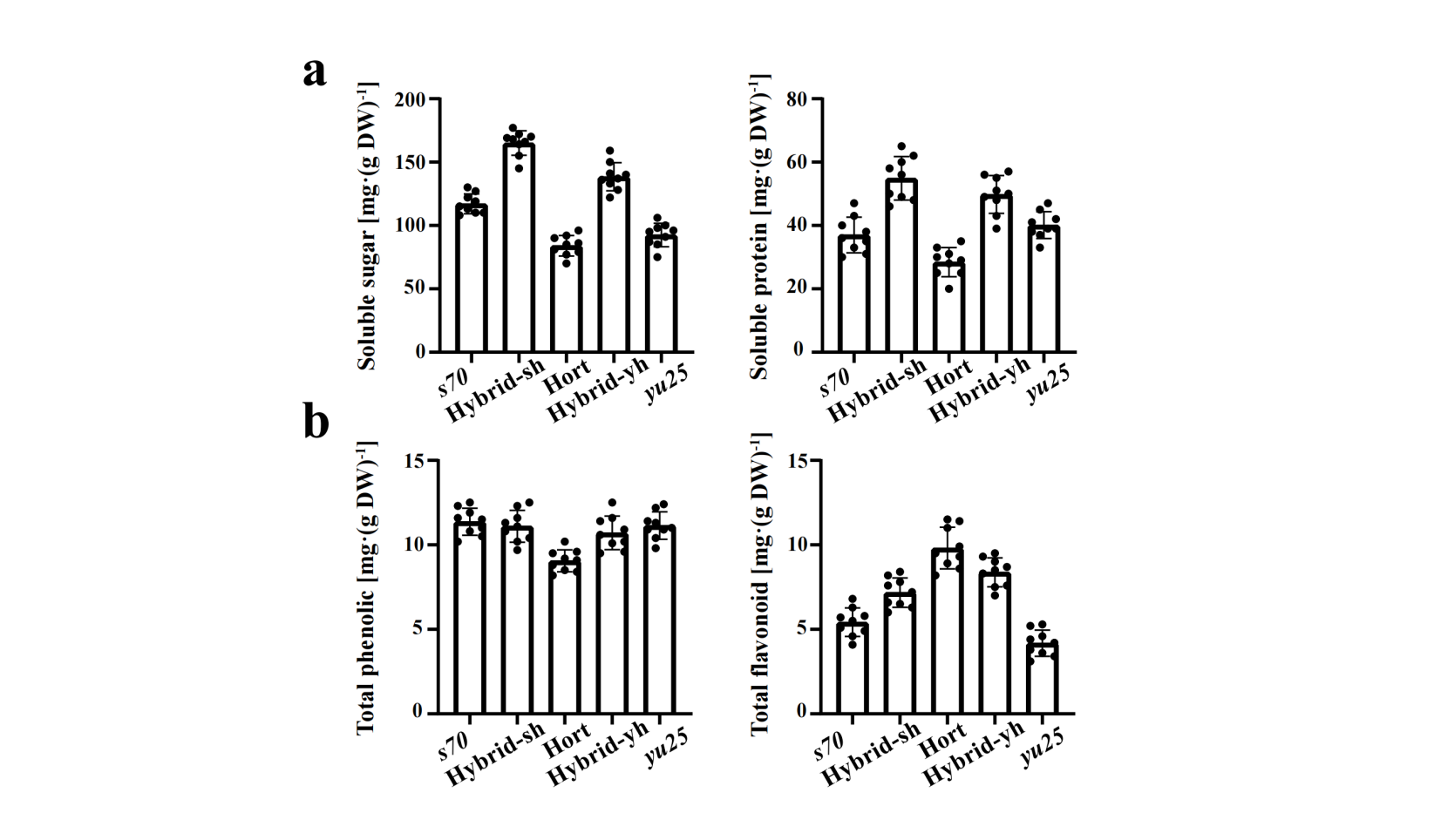

**Figure S1. Determination of primary metabolites.**

The bar graph showed the content of soluble sugars and proteins **(a)**, total phenols, and flavonoids **(b)** in F_1_ hybrids and their relative parents.

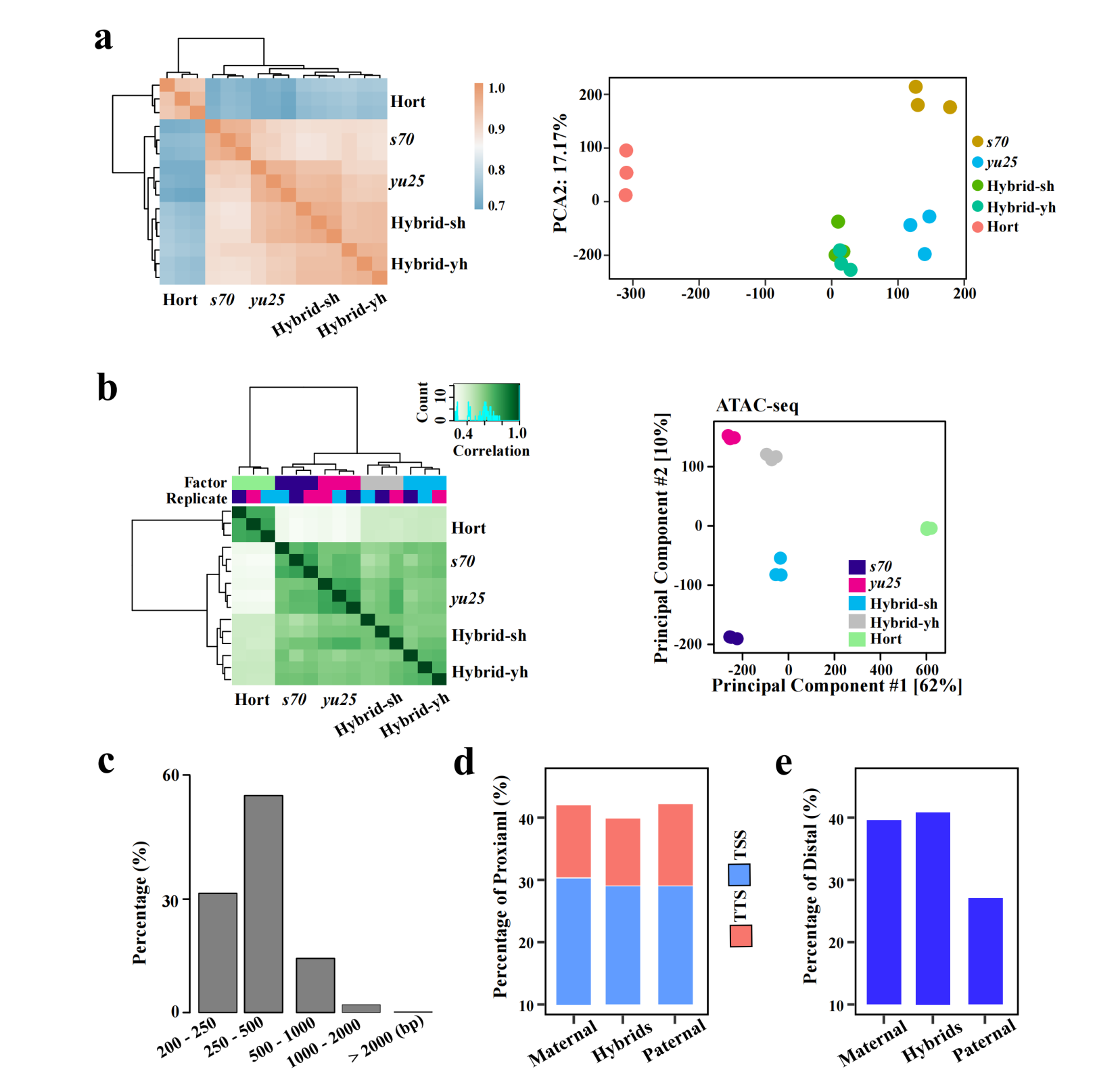

**Figure S2. Quality control of ATAC-seq and RNA-seq datasets.**

PCA plots (left) and Spearman's rank correlation coefficient heatmap (right) of RNA-seq **(a)** and ATAC-seq **(b)**. **(c)** The bar graph showed the percentage of ACRs with different lengths (bp). **(d)** - **(e)**, The bar graph showed the percentage of Proximal (including TSS and TTS) ACRs **(d)**, and Distal ACRs **(e)** in the hybrids and their parents.

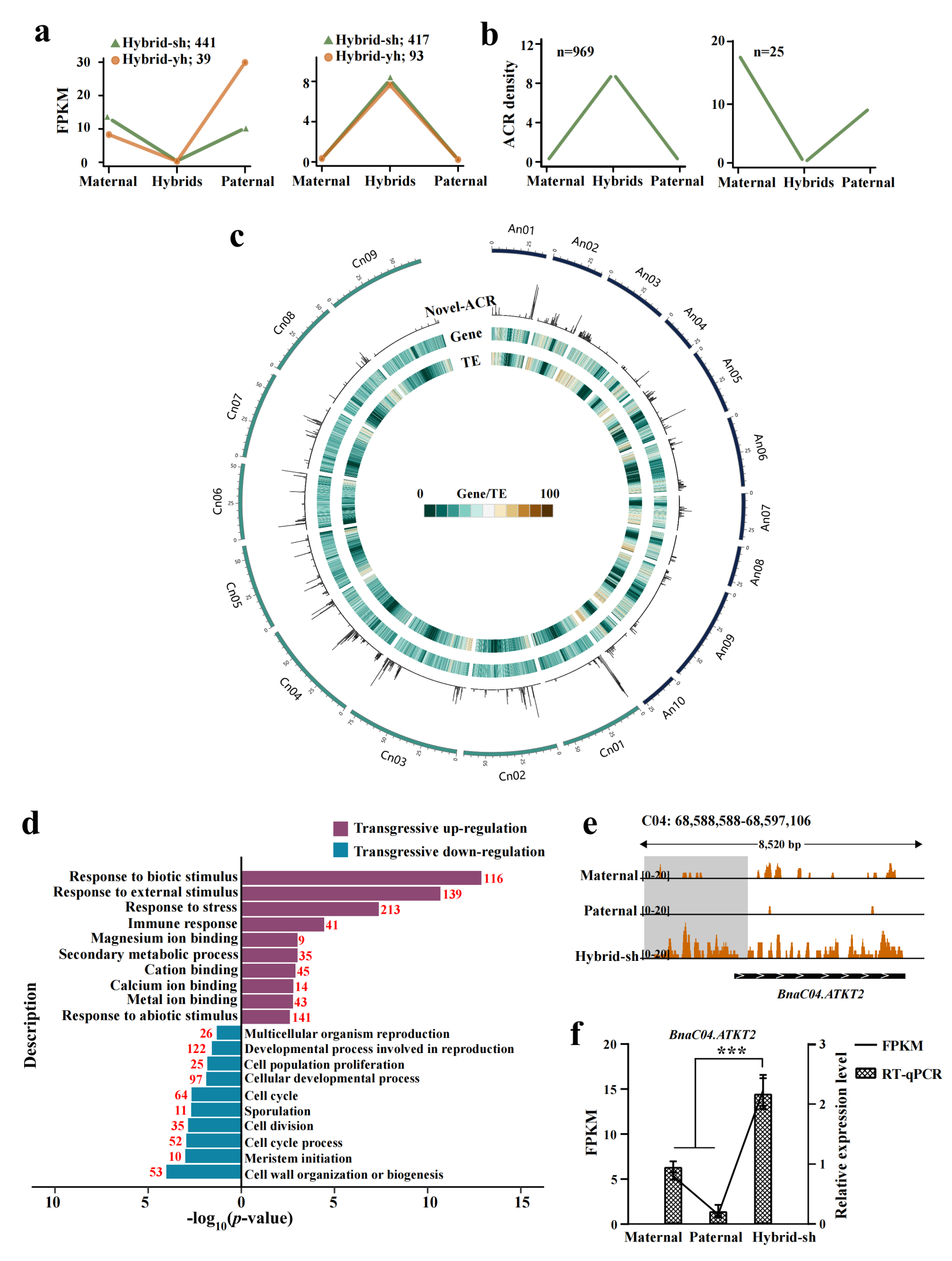

**Figure S3. Genome-wide distribution of different types of ACRs in F_1_ hybrids.**

**(a)** The graph showed the number of genes with novel expressed and silenced expressed patterns in the F_1_ hybrids. The genes with FPKM > 1 in the F_1_ hybrids and FPKM < 0.1 in two parents were defined as novel expressed genes. The genes with FPKM < 0.1 in the F_1_ hybrids and FPKM < 1 in two parents were defined as silenced expressed genes. **(b)** The graph showed the ACR density of novel and silenced ACRs in the F_1_ hybrids. The ACRs with Reads > 5 in the F_1_ hybrids and Read = 0 in two parents were defined as novel expressed ACRs. The ACRs with Reads = 0 in the F_1_ hybrids and Reads > 5 in two parents were defined as silenced expressed ACRs. **(c)** Browser Circos plots showed the genome distribution of genes, TEs, and novel-ACR regions in the F_1_ hybrids. **(d)** GO enrichment analyses of transgressive regulation genes. The top 10 biological process GO terms were listed. **(e)** Genome browser showed ATAC-seq peaks around *Potassium Transporter 2* (*POT2*) in the Hybrid-sh and its relative parents. **(f)** The graph showed the gene expression of *BnaC04.POT2* in Hybrid-sh and its relative parents. Error bars indicated the mean ± SD of three biological replicates. The Student’s t-test; ****p* < 0.001.

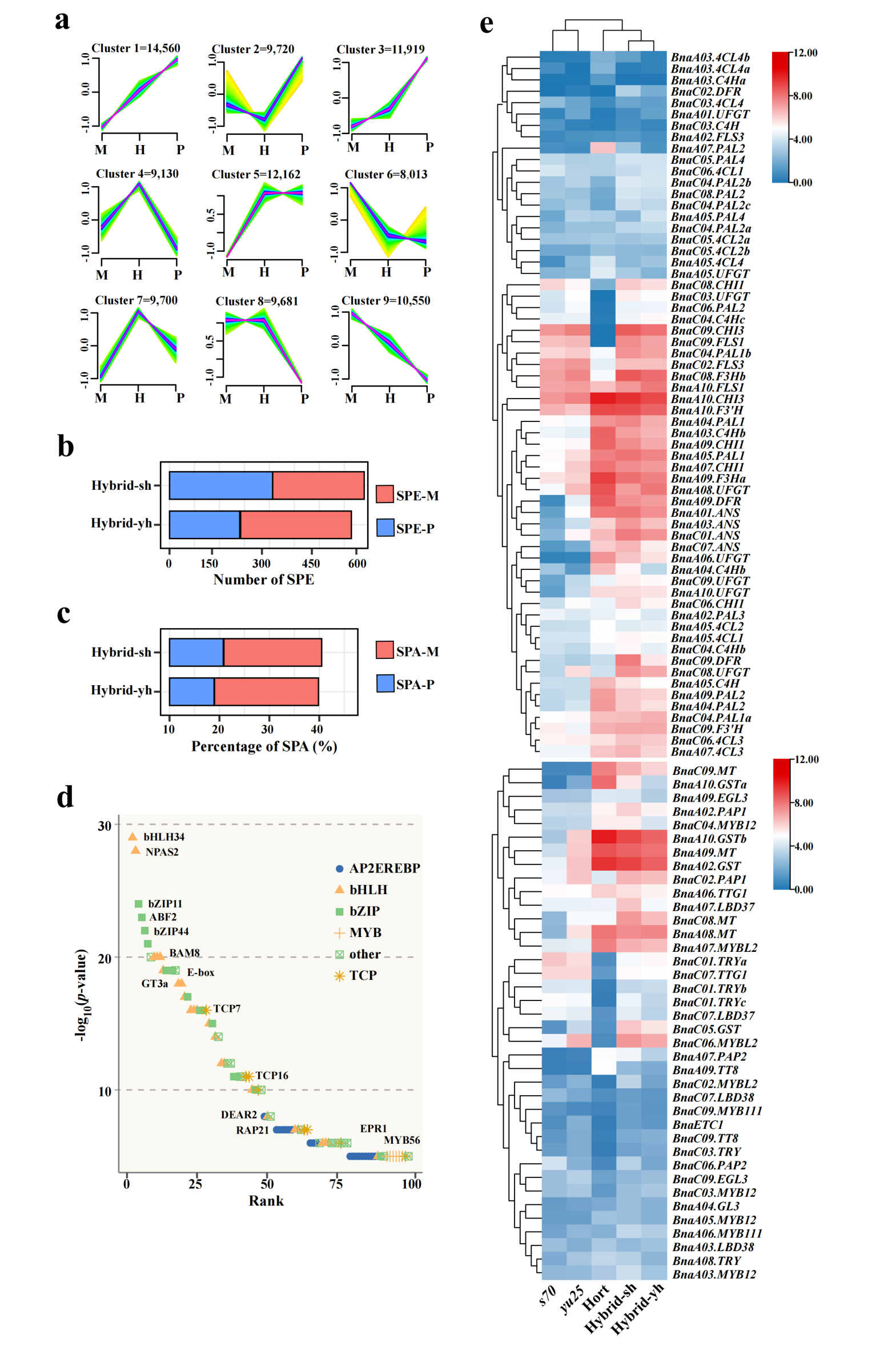

**Figure S4. Characterization of SPA-ACRs in the F_1_ hybrids.**

**(a)** The graphs showed the *c*-means soft clustering analysis of chromatin accessibility levels of A subgeomes in the Hybrid-yh and its relative parents. **(b)** The graph showed the number of SPE-M and SPE-P genes in the A subgeomes of Hybird-sh and Hybird-yh. **(c)** The graph showed the percentage of SPA-M and SPA-P genes in the A subgeomes of Hybird-sh and Hybird-yh. **(d)** The graph showed the ranking of enriched motifs in Hybrid-sh ACRs. Colored dots represented transcription factors. The *p*-value for each subject was estimated using AME software. **(e)** The heatmap showed the expression levels of anthocyanin biosynthesis genes and the related transcription factors in the hybrids and their parents. The colored boxes represent the gene expression heatmap, normalized by log_2_FPKM values obtained from RNA-seq.

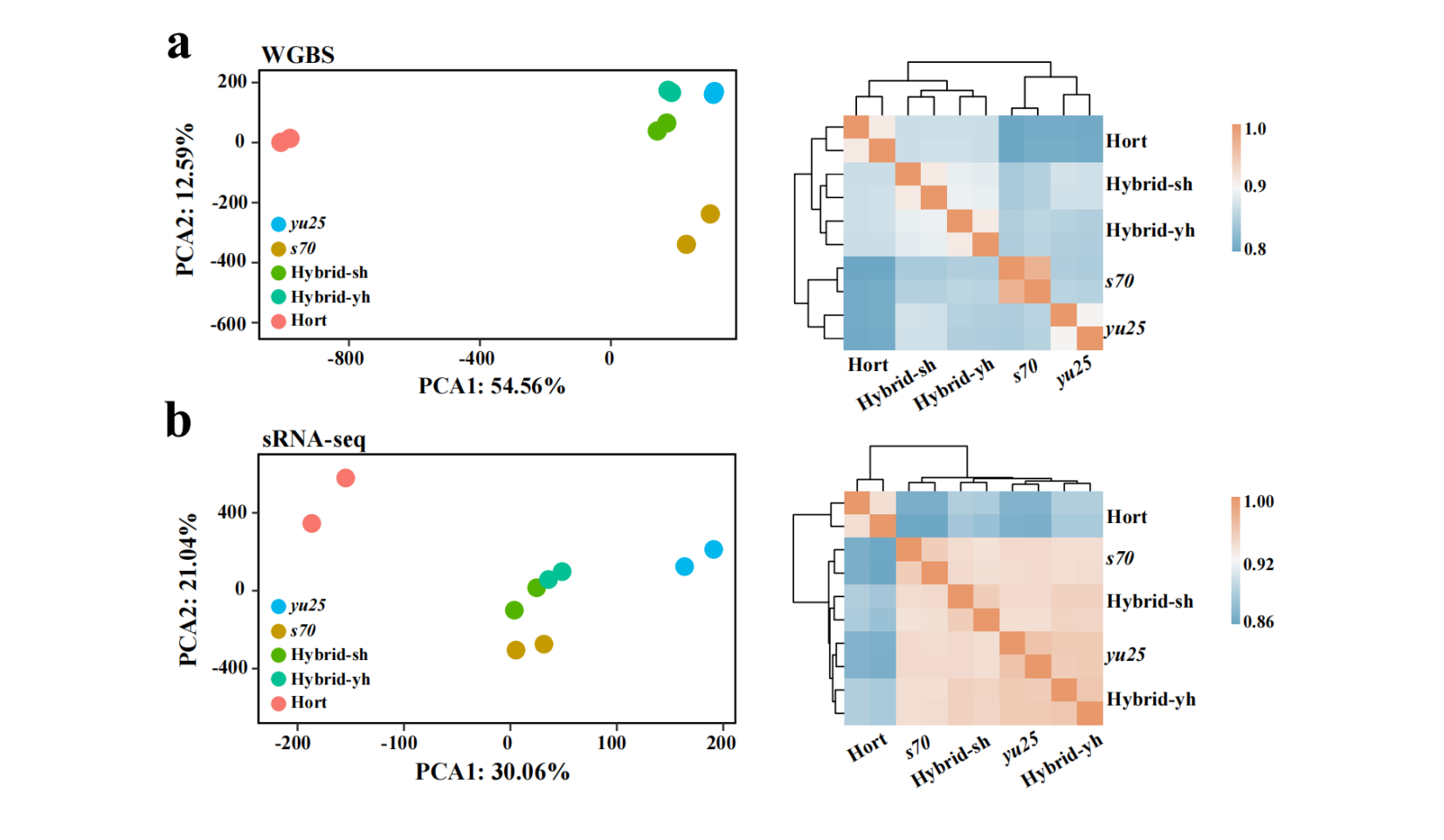

**Figure S5. Quality control of WGBS and sRNA-seq datasets.**

The graphs showed the PCA plots (left) and Spearman's rank correlation coefficients (right) of WGBS **(a)** and sRNA-seq **(b)** datasets.

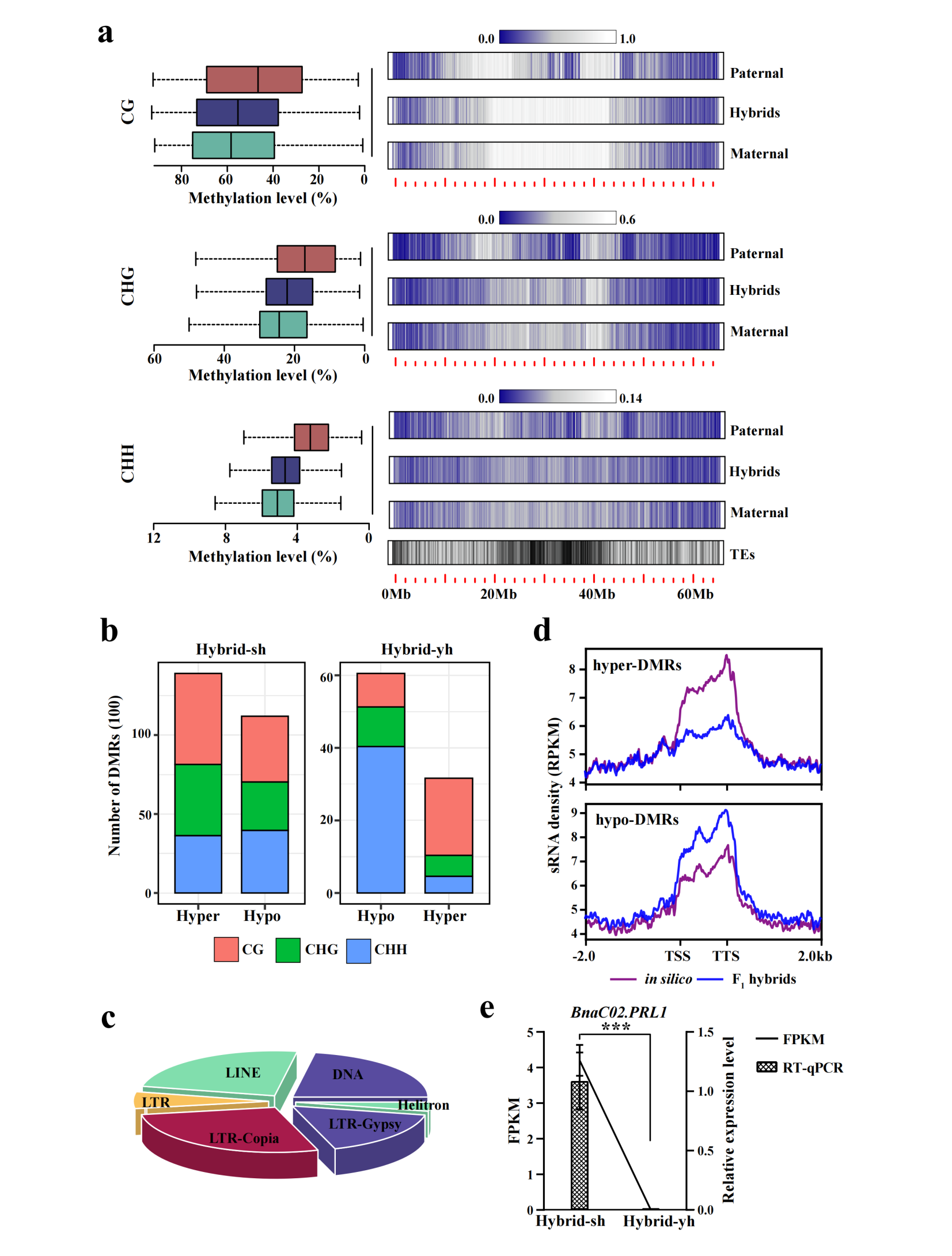

**Figure S6. Characterization of hyper- and hypo-DMRs in the F_1_ hybrids.**

**(a)** The boxplots showed the total DNA methylation level (CG, CHG, and CHH) of F_1_ hybrids (Hybird-sh (left) and Hybird-yh (right)) and *in silico* hybrids (Wilcoxon rank-sum test; ***p* < 0.01; ****p* < 0.001; ns, no significant difference). **(b)** The bar graph showed the number of hyper- and hypo-DMRs in DNA methylation (CG, CHG, and CHH contexts) in Hybrid-sh and Hybrid-yh. **(c)** The pie chart showed the proportion of TE-derived MDRs (TE-driven DMRs when more than 50% of the region overlaps with TEs). **(c)** The bar graph showed the percentage of TE in TCM and TCdM. **(d)** The graph showed the sRNA densities of hyper-DMRs (top) and hypo-DMRs (bottom). **(e)** The graph showed the gene expression of *BnaC02.PRL1* in the Hybrid-sh and Hybrid-yh plants. Error bars indicated the mean ± SD of three biological replicates. The Student’s *t*-test; ****p* < 0.001.

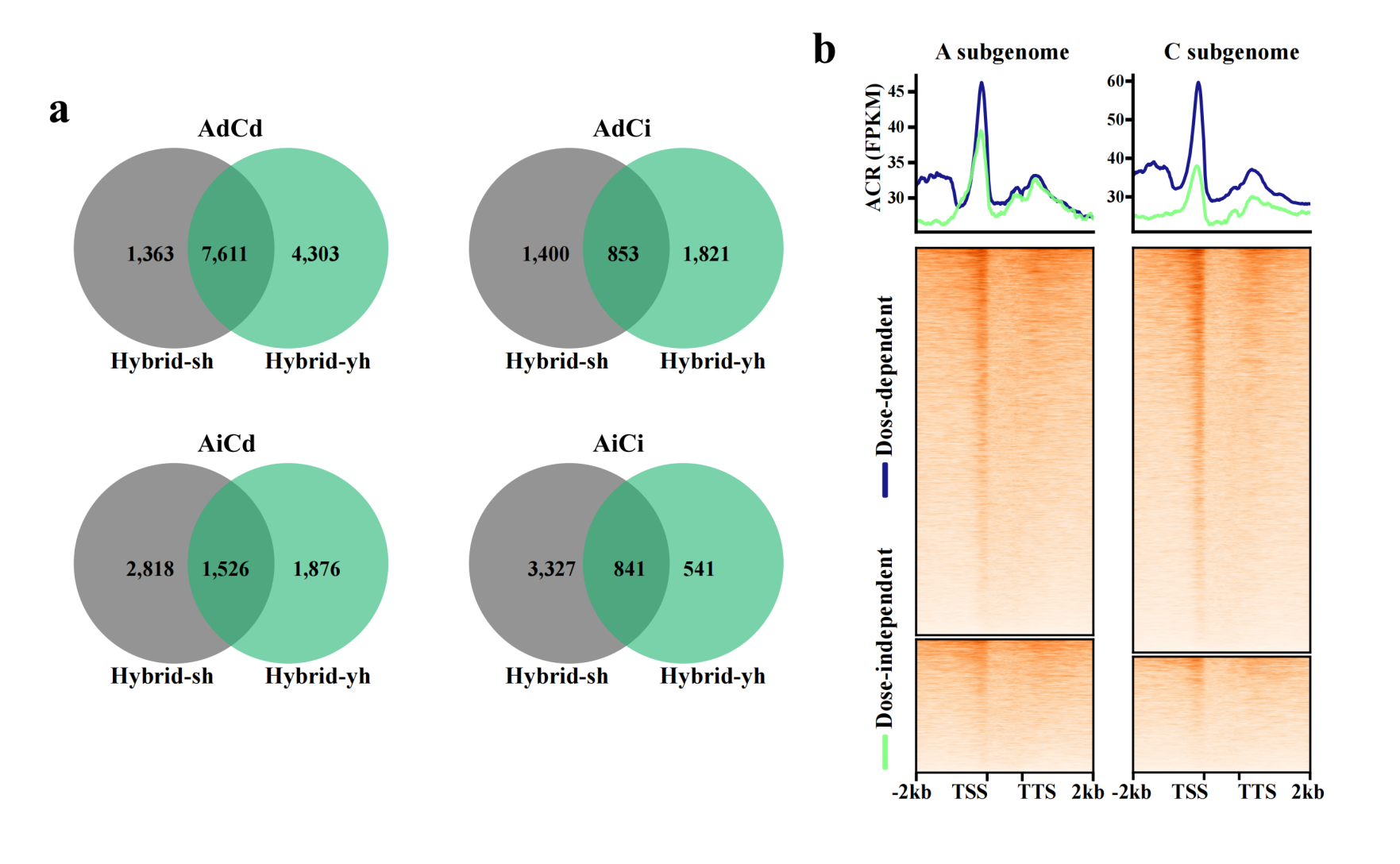

**Figure S7. Dose-dependent and dose-independent distributions in the A and C subgenomes.**

**(a)** Venn diagram showed the number of the overlapping AdCd, AdCi, AiCd and AiCi genes in Hybrid-sh and Hybrid-yh. **(b)** The graph showed the dose-dependent and dose-independent ACR density distribution in the A and C subgenomes.

**Supplemental Table S1. Primer sequences used for RT-qPCR analyses.**

| **Gene_id** | **Forward primer (5'-3')** | **Reverse primer (5'-3')** |
| --- | --- | --- |
| *BnaA07.PAP2* | GCTGGTCTAAATCGGTGCAG | CGACCGGGTAATCTACCAGC |
| *BnaC02.PRL1* | GCTCTTGCATTCCCAGGTT | ACGATCCACTGCTTTTGCTAC |
| *BnaA06.UF3GT* | GGTTTTCTACGATTCCGCTGA | TTCTTCCTCACTAGAGGCGG |
| *BnaA01.POT4* | CTCAGCTTTCATGGGCGAG | AGCGTGAGGGTCCAGAAAATC |
| *BnaC04.POT2* | CAGGCAAGTCGCTGATGAG | ACAGCAGAGAACACGGAGATAG |
| *Actin7* | GGCATCACACTTTCTACAACGA | GCACAATACCAGTAGTACGTCC |

**Supplemental Table S2. Statistics of RNA-seq data and reads mapping for all samples.**

| **RNA-seq** | **Raw Reads** | **Clean Reads** | **Q30(%)** | **GC Content (%)** | **Mapped Clean reads** |
| --- | --- | --- | --- | --- | --- |
| **s70** | 20674100 | 19215459 | 93.65 | 47.31 | 1802602209 |
| **s70** | 20541951 | 19201504 | 93.82 | 47.23 | 1807821602 |
| **s70** | 21956773 | 20827026 | 93.57 | 46.45 | 1957115633 |
| **yu25** | 21721793 | 20540690 | 93.22 | 47.51 | 1937808695 |
| **yu25** | 21370948 | 20133806 | 93.05 | 47.2 | 1899423258 |
| **yu25** | 21011896 | 20546666 | 93.39 | 46.7 | 1955631670 |
| **Hybrid-sh** | 21487410 | 20941988 | 92.71 | 46.78 | 1984043943 |
| **Hybrid-sh** | 21434438 | 20589841 | 93.19 | 46.38 | 1964888527 |
| **Hybrid-sh** | 19786691 | 18711457 | 93.78 | 46.81 | 1765800197 |
| **Hybrid-yh** | 19760029 | 18592257 | 93.19 | 46.86 | 1764219267 |
| **Hybrid-yh** | 19900711 | 18869088 | 93.09 | 47.3 | 1782374052 |
| **Hybrid-yh** | 21251984 | 20003638 | 93.38 | 47.19 | 1898545283 |
| **hort** | 21862561 | 20822153 | 93.21 | 47.27 | 1850048294 |
| **hort** | 22985725 | 21883259 | 93.65 | 47.3 | 1946953553 |
| **hort** | 21643208 | 20421458 | 93.59 | 47.35 | 1810158037 |

**Supplemental Table S3. Statistics of ATAC-seq data and reads mapping for all samples.**

| **ATAC-seq** | **Raw Reads** | **Clean Reads** | **Q30(%)** | **SPOT** | **Mapped Clean reads** |
| --- | --- | --- | --- | --- | --- |
| **s70** | 46713984 | 46415584 | 94.46 | 0.28 | 41924249 |
| **s70** | 48946191 | 48635311 | 92.02 | 0.26 | 37994096 |
| **s70** | 50592226 | 50294108 | 92.2 | 0.29 | 39556686 |
| **yu25** | 51058619 | 50755263 | 92.48 | 0.28 | 38861401 |
| **yu25** | 49497213 | 49216199 | 92.46 | 0.26 | 38194516 |
| **yu25** | 48241753 | 47956390 | 92.01 | 0.28 | 37652044 |
| **Hybrid-sh** | 44186451 | 43851231 | 91.57 | 0.28 | 31213248 |
| **Hybrid-sh** | 50010924 | 49605541 | 92.27 | 0.28 | 38723274 |
| **Hybrid-sh** | 50513122 | 50120282 | 92.34 | 0.26 | 38399968 |
| **Hybrid-yh** | 45181181 | 44702269 | 92.31 | 0.25 | 33388099 |
| **Hybrid-yh** | 48940370 | 48488343 | 91.77 | 0.28 | 38499425 |
| **Hybrid-yh** | 41344897 | 41029470 | 92.07 | 0.25 | 29966350 |
| **hort** | 45155912 | 44639786 | 90.71 | 0.32 | 35038745 |
| **hort** | 48043231 | 47614365 | 91.28 | 0.32 | 36007670 |
| **hort** | 44909946 | 44517771 | 91.23 | 0.33 | 34211196 |

**Supplemental Table S4. GO functional categories of non-additively up-regulated genes in the Hybrid-sh.**

| **GO_ID** | ***P*_value** | **Count** | **Description** |
| --- | --- | --- | --- |
| GO:0009812 | 1.56E-04 | 23 | flavonoid metabolic process |
| GO:0009813 | 6.84E-04 | 18 | flavonoid biosynthetic process |
| GO:0051553 | 9.60E-04 | 7 | flavone biosynthetic process |
| GO:0009607 | 3.87E-14 | 176 | response to biotic stimulus |
| GO:0050896 | 5.67E-13 | 599 | response to stimulus |
| GO:0009605 | 4.64E-12 | 217 | response to external stimulus |
| GO:0009409 | 2.42E-10 | 82 | response to cold |
| GO:0009725 | 1.21E-07 | 220 | response to hormone |
| GO:0009719 | 2.18E-07 | 225 | response to endogenous stimulus |
| GO:0006629 | 8.45E-06 | 101 | lipid metabolic process |
| GO:0006631 | 3.11E-05 | 37 | fatty acid metabolic process |
| GO:0046283 | 1.02E-04 | 15 | anthocyanin-containing compound metabolic process |
| GO:0009733 | 1.29E-04 | 54 | response to auxin |
| GO:0044042 | 4.10E-04 | 29 | glucan metabolic process |
| GO:0051552 | 1.32E-03 | 7 | flavone metabolic process |

**Supplemental Table S5. GO functional categories of non-additively up-regulated genes in the Hybrid-yh.**

| **GO_ID** | ***P*_value** | **Count** | **Description** |
| --- | --- | --- | --- |
| GO:0042218 | 7.89E-08 | 5 | 1-aminocyclopropane-1-carboxylate biosynthetic process |
| GO:0010025 | 1.72E-05 | 6 | wax biosynthetic process |
| GO:1901570 | 3.56E-05 | 6 | fatty acid derivative biosynthetic process |
| GO:0009733 | 1.67E-03 | 15 | response to auxin |
| GO:0044255 | 2.94E-03 | 22 | cellular lipid metabolic process |
| GO:0072330 | 3.63E-03 | 11 | monocarboxylic acid biosynthetic process |
| GO:0009741 | 4.28E-03 | 7 | response to brassinosteroid |
| GO:0006629 | 4.80E-03 | 23 | lipid metabolic process |
| GO:0015698 | 8.35E-03 | 7 | inorganic anion transport |
| GO:0009812 | 1.46E-03 | 19 | flavonoid metabolic process |
| GO:0046283 | 4.14E-05 | 15 | anthocyanin biosynthesis process. |
| GO:0044042 | 3.16E-04 | 22 | glucan biosyntheti process |

**Supplemental Table S6. The contents of 16 ionic components in the F_1_ hybrids and their parents.**

|  | **s70** | **yu25** | **Hybrid**  **-sh** | **Hybrid**  **-yh** | **hort** | **Mean** | **SD^1^** | **CV^1^** |
| --- | --- | --- | --- | --- | --- | --- | --- | --- |
| Mg^2^ | 2.61 | 2.96 | 4.79 | 2.77 | 2.09 | 3.04 | 1.03 | 33.84 |
| P | 7.86 | 6.05 | 10.58 | 8.42 | 5.04 | 7.59 | 2.16 | 28.41 |
| K | 22.01 | 27.98 | 45.25 | 36.07 | 34.23 | 30.91 | 8.00 | 25.89 |
| Ca | 8.04 | 6.99 | 14.77 | 9.66 | 6.29 | 9.15 | 3.39 | 37.04 |
| Na | 1.88 | 1.95 | 1.27 | 1.14 | 1.67 | 1.58 | 0.36 | 22.83 |
| B^2^ | 31.22 | 39.41 | 28.57 | 25.91 | 32.47 | 31.52 | 5.09 | 16.14 |
| Se | 0.10 | 0.09 | 0.11 | 0.10 | 0.07 | 0.09 | 0.02 | 17.21 |
| Mn | 20.15 | 20.02 | 21.80 | 19.63 | 18.13 | 19.95 | 1.31 | 6.58 |
| Fe | 46.69 | 51.37 | 46.50 | 46.51 | 62.83 | 50.78 | 7.05 | 13.89 |
| Cu | 3.09 | 3.45 | 3.66 | 3.74 | 5.11 | 3.81 | 0.77 | 20.25 |
| Zn | 49.65 | 53.53 | 43.91 | 41.98 | 47.25 | 47.26 | 4.59 | 9.7 |
| Mo | 0.30 | 0.38 | 0.35 | 0.36 | 0.49 | 0.37 | 0.07 | 18.21 |
| Ba | 10.80 | 11.75 | 7.67 | 8.03 | 9.54 | 9.56 | 1.75 | 18.3 |
| As | 0.03 | 0.05 | 0.03 | 0.02 | 0.01 | 0.03 | 0.02 | 59.47 |
| Cd | 0.05 | 0.08 | 0.12 | 0.11 | 0.26 | 0.12 | 0.08 | 64.14 |
| Pb | 0.01 | 0.01 | 0.01 | 0.01 | 0.01 | 0.01 | 0.00 | 15.12 |

1. The parameters shown are: standard deviation (± SD from five replicates), Coefficient of Variation (CV, %).
2. The units of 5 macroelements (Mg, P, K, Ca, and Na) are (g·kg-1), and the units of 11 trace elements (B, Se, Mn, Fe, Cu, Zn, Mo, Ba, As, Cd, and Pb) are (mg·kg-1).

**Supplemental Table S7. Statistics of WGBS-seq data and reads mapping for all samples.**

| **WGBS** | **QC-passed reads** | **Properly paired reads** | **Average Cytosine depth** | **Bisulfite conversion (%)** | **mC/(C+T) (%)** |
| --- | --- | --- | --- | --- | --- |
| **s70** | 203755516 | 192649161 | 30.3139 | 99.58 | 10.61 |
| **s70** | 242642051 | 196800471 | 33.6186 | 99.21 | 13.29 |
| **yu25** | 203750557 | 193562163 | 30.2654 | 99.32 | 14.47 |
| **yu25** | 193436940 | 184328420 | 28.8405 | 99.45 | 12.95 |
| **Hybrid-sh** | 199423059 | 188154639 | 29.6604 | 99.26 | 12.52 |
| **Hybrid-sh** | 206212634 | 195152714 | 30.6538 | 99.38 | 10.26 |
| **Hybrid-yh** | 184643639 | 173473362 | 27.416 | 99.41 | 12.03 |
| **Hybrid-yh** | 207849273 | 197326844 | 30.8683 | 99.55 | 11.03 |
| **hort** | 193809351 | 177983477 | 28.5439 | 99.16 | 9.85 |
| **hort** | 238282559 | 219909569 | 35.0476 | 99.23 | 10.42 |

**Supplemental Table S8. Statistics of sRNA-seq data and reads mapping for all samples.**

| **sRNA-seq** | **Raw Reads** | **Clean Reads** | **Q20 (%)** | **Q30 (%)** | **Mapped Clean reads** |
| --- | --- | --- | --- | --- | --- |
| **s70** | 29806536 | 28640685 | 99.3 | 97.7 | 26041698 |
| **s70** | 28599807 | 27126278 | 99.1 | 97.1 | 26482857 |
| **yu25** | 29752067 | 28267018 | 99.1 | 96.9 | 27275545 |
| **yu25** | 30151921 | 28854786 | 98.2 | 95.1 | 27904231 |
| **Hybrid-sh** | 29158016 | 27039422 | 98.4 | 95.6 | 26166560 |
| **Hybrid-sh** | 28095774 | 27315576 | 98.6 | 95.9 | 26116061 |
| **Hybrid-yh** | 28393134 | 27749028 | 98.5 | 95.7 | 26825736 |
| **Hybrid-yh** | 28179167 | 27519684 | 98.2 | 95 | 26567866 |
| **hort** | 28700923 | 28020472 | 98.9 | 96.5 | 26911249 |
| **hort** | 28403184 | 27835112 | 99.1 | 97.1 | 26416568 |
